## Supplemental codes for "Detection, Isolation and Quantification of Myocardial Infarct with Four Different Histological Staining Techniques": fibrosis_analyzer.m

```
function varargout = fibrosis_analyzer(varargin)
% FIBROSIS_ANALYZER MATLAB code for fibrosis_analyzer.fig
%      FIBROSIS_ANALYZER, by itself, creates a new FIBROSIS_ANALYZER or raises the existing
%      singleton*.
%
%      H = FIBROSIS_ANALYZER returns the handle to a new FIBROSIS_ANALYZER or the handle to
%      the existing singleton*.
%
%      FIBROSIS_ANALYZER('CALLBACK',hObject,eventData,handles,...) calls the local
%      function named CALLBACK in FIBROSIS_ANALYZER.M with the given input arguments.
%
%      FIBROSIS_ANALYZER('Property','Value',...) creates a new FIBROSIS_ANALYZER or raises the
%      existing singleton*.  Starting from the left, property value pairs are
%      applied to the GUI before fibrosis_analyzer_OpeningFcn gets called.  An
%      unrecognized property name or invalid value makes property application
%      stop.  All inputs are passed to fibrosis_analyzer_OpeningFcn via varargin.
%
%      *See GUI Options on GUIDE's Tools menu.  Choose "GUI allows only one
%      instance to run (singleton)".
%
% See also: GUIDE, GUIDATA, GUIHANDLES

% Edit the above text to modify the response to help fibrosis_analyzer

% Last Modified by GUIDE v2.5 16-Jul-2024 11:11:24

% Begin initialization code - DO NOT EDIT
gui_Singleton = 1;
gui_State = struct('gui_Name',       mfilename, ...
                   'gui_Singleton',  gui_Singleton, ...
                   'gui_OpeningFcn', @fibrosis_analyzer_OpeningFcn, ...
                   'gui_OutputFcn',  @fibrosis_analyzer_OutputFcn, ...
                   'gui_LayoutFcn',  [] , ...
                   'gui_Callback',   []);
if nargin && ischar(varargin{1})
    gui_State.gui_Callback = str2func(varargin{1});
end

if nargout
    [varargout{1:nargout}] = gui_mainfcn(gui_State, varargin{:});
else
    gui_mainfcn(gui_State, varargin{:});
end
% End initialization code - DO NOT EDIT


% --- Executes just before fibrosis_analyzer is made visible.
function fibrosis_analyzer_OpeningFcn(hObject, eventdata, handles, varargin)
% This function has no output args, see OutputFcn.
% hObject    handle to figure
% eventdata  reserved - to be defined in a future version of MATLAB
% handles    structure with handles and user data (see GUIDATA)
% varargin   command line arguments to fibrosis_analyzer (see VARARGIN)

% Choose default command line output for fibrosis_analyzer
handles.output = hObject;

flg_data_folder=0;
bg_indicator=0;
handles.flg_data_folder=flg_data_folder;
handles.bg_indicator=bg_indicator;
% Update handles structure
guidata(hObject, handles);

% UIWAIT makes fibrosis_analyzer wait for user response (see UIRESUME)
% uiwait(handles.figure1);


% --- Outputs from this function are returned to the command line.
function varargout = fibrosis_analyzer_OutputFcn(hObject, eventdata, handles) 
% varargout  cell array for returning output args (see VARARGOUT);
% hObject    handle to figure
% eventdata  reserved - to be defined in a future version of MATLAB
% handles    structure with handles and user data (see GUIDATA)

% Get default command line output from handles structure
varargout{1} = handles.output;


% --- Executes on button press in load_image.
function load_image_Callback(hObject, eventdata, handles)
% hObject    handle to load_image (see GCBO)
% eventdata  reserved - to be defined in a future version of MATLAB
% handles    structure with handles and user data (see GUIDATA)

set(handles.extraction_thd,'String', '1.2');
set(handles.tissue_pixels,'String', ' ');
set(handles.fibrosis_pixels,'String', ' ');
set(handles.fibrosis_percent,'String', ' ');
set(handles.tissue_thd_black,'String', '0.2');
set(handles.tissue_thd_white,'String', '0.8');

set(handles.save_data,'ForegroundColor','k');


[filname,pathname] = uigetfile({'*.tif';'*.png';'*.jpg';'*.gif'}, 'Select File to work with');
img_ori = imread([pathname, '\', filname]);

set(handles.file_name_display,'String',filname);


cla(handles.axes1);
cla(handles.axes2);
cla(handles.axes3);
reset(handles.axes1);
reset(handles.axes2);
reset(handles.axes3);

axes(handles.axes1)
imshow(img_ori)
title('Original Image');
axis off

handles.img_ori=img_ori;
handles.fname_ori=filname;
handles.pname_ori=pathname;

guidata(hObject, handles);


function extraction_thd_Callback(hObject, eventdata, handles)
% hObject    handle to extraction_thd (see GCBO)
% eventdata  reserved - to be defined in a future version of MATLAB
% handles    structure with handles and user data (see GUIDATA)

% Hints: get(hObject,'String') returns contents of extraction_thd as text
%        str2double(get(hObject,'String')) returns contents of extraction_thd as a double


% --- Executes during object creation, after setting all properties.
function extraction_thd_CreateFcn(hObject, eventdata, handles)
% hObject    handle to extraction_thd (see GCBO)
% eventdata  reserved - to be defined in a future version of MATLAB
% handles    empty - handles not created until after all CreateFcns called

% Hint: edit controls usually have a white background on Windows.
%       See ISPC and COMPUTER.
if ispc && isequal(get(hObject,'BackgroundColor'), get(0,'defaultUicontrolBackgroundColor'))
    set(hObject,'BackgroundColor','white');
end


% --- Executes on button press in extract_fibrosis.
function extract_fibrosis_Callback(hObject, eventdata, handles)
% hObject    handle to extract_fibrosis (see GCBO)
% eventdata  reserved - to be defined in a future version of MATLAB
% handles    structure with handles and user data (see GUIDATA)
set(handles.fibrosis_pixels,'String', ' ');
img_ori=handles.img_ori;
img_double=im2double(img_ori);

set(handles.fibrosis_pixels,'String', ' ');
con_thd=str2num(get(handles.extraction_thd,'string'));

[row,col,clr]=size(img_double);
img_thd=zeros(row,col,clr);
pixels_fb=0;
for b=1:1:row
        for c=1:1:col
            if img_double(b,c,1)<0.3&&img_double(b,c,2)<0.3&&img_double(b,c,3)<0.3
             img_thd(b,c,:)=1;
            elseif img_double(b,c,1)>0.8&&img_double(b,c,2)>0.8&&img_double(b,c,3)>0.8
             img_thd(b,c,:)=1;
            else
                if img_double(b,c,3)/img_double(b,c,1)>=con_thd
                img_thd(b,c,1)=0;
                img_thd(b,c,2)=0;
                img_thd(b,c,3)=img_double(b,c,3);
                pixels_fb=pixels_fb+1;
                elseif img_double(b,c,3)/img_double(b,c,1)<con_thd
                 img_thd(b,c,1)=1;
                img_thd(b,c,2)=1;
                img_thd(b,c,3)=1;
                end
            end
        end
        
         
end


% h=fspecial('disk',5);
 img_filtered=img_thd;

axes(handles.axes2)
imshow(img_filtered)
title('Extracted Fibrosis')
axis off

% pixels_fb=0;
% for b=1:1:row
%       for c=1:1:col
%           if img_filtered(b,c,3)~=0
%                 pixels_fb=pixels_fb+1;
%           end
%       end
% end

set(handles.fibrosis_pixels,'String', num2str(pixels_fb));

if get(handles.tissue_pixels,'String')==' '
    set(handles.fibrosis_percent,'String', ' ');
else 
    fibrosis_percent=pixels_fb*100/str2num(get(handles.tissue_pixels,'String'));
    fibrosis_percent=round(fibrosis_percent,2);
    set(handles.fibrosis_percent,'String',num2str(fibrosis_percent));
end

handles.img_highlighted=img_filtered;
handles.row_img=row;
handles.col_img=col;

guidata(hObject, handles);


% --- Executes on button press in extract_tissue.
function extract_tissue_Callback(hObject, eventdata, handles)
% hObject    handle to extract_tissue (see GCBO)
% eventdata  reserved - to be defined in a future version of MATLAB
% handles    structure with handles and user data (see GUIDATA)
img_ori=handles.img_ori;
img_double=im2double(img_ori);

set(handles.tissue_pixels,'String', ' ');
[row,col,clr]=size(img_double);
img_thd=zeros(row,col,clr);
black_thd=str2num(get(handles.tissue_thd_black,'String'));
white_thd=str2num(get(handles.tissue_thd_white,'String'));
pixels_tissue=0;
for b=1:1:row
        for c=1:1:col
            
            % remove the black background
           if img_double(b,c,1)<black_thd&&img_double(b,c,2)<black_thd&&img_double(b,c,3)<black_thd
               img_thd(b,c,1)=1;
               img_thd(b,c,2)=1;
               img_thd(b,c,3)=1;
           elseif img_double(b,c,1)>white_thd&&img_double(b,c,2)>white_thd&&img_double(b,c,3)>white_thd
               img_thd(b,c,1)=1;
               img_thd(b,c,2)=1;
               img_thd(b,c,3)=1;
           else
               img_thd(b,c,:)=img_double(b,c,:);
               pixels_tissue=pixels_tissue+1;
           end
        end
         
end

axes(handles.axes3)
imshow(img_thd)
title('Extracted Tissue')
axis off

% pixels_tissue=0;
% for b=1:1:row
%       for c=1:1:col
%           if img_thd(b,c,3)~=0
%                 pixels_tissue=pixels_tissue+1;
%           end
%       end
% end

set(handles.tissue_pixels,'String', num2str(pixels_tissue));


if get(handles.fibrosis_pixels,'String')==' '
    set(handles.fibrosis_percent,'String', ' ');
else 
    fibrosis_percent=str2num(get(handles.fibrosis_pixels,'String'))*100/pixels_tissue;
    fibrosis_percent=round(fibrosis_percent,2);
    set(handles.fibrosis_percent,'String',num2str(fibrosis_percent));
end


handles.img_tissue=img_thd;
guidata(hObject, handles)

function tissue_pixels_Callback(hObject, eventdata, handles)
% hObject    handle to tissue_pixels (see GCBO)
% eventdata  reserved - to be defined in a future version of MATLAB
% handles    structure with handles and user data (see GUIDATA)

% Hints: get(hObject,'String') returns contents of tissue_pixels as text
%        str2double(get(hObject,'String')) returns contents of tissue_pixels as a double


% --- Executes during object creation, after setting all properties.
function tissue_pixels_CreateFcn(hObject, eventdata, handles)
% hObject    handle to tissue_pixels (see GCBO)
% eventdata  reserved - to be defined in a future version of MATLAB
% handles    empty - handles not created until after all CreateFcns called

% Hint: edit controls usually have a white background on Windows.
%       See ISPC and COMPUTER.
if ispc && isequal(get(hObject,'BackgroundColor'), get(0,'defaultUicontrolBackgroundColor'))
    set(hObject,'BackgroundColor','white');
end


function fibrosis_pixels_Callback(hObject, eventdata, handles)
% hObject    handle to fibrosis_pixels (see GCBO)
% eventdata  reserved - to be defined in a future version of MATLAB
% handles    structure with handles and user data (see GUIDATA)

% Hints: get(hObject,'String') returns contents of fibrosis_pixels as text
%        str2double(get(hObject,'String')) returns contents of fibrosis_pixels as a double


% --- Executes during object creation, after setting all properties.
function fibrosis_pixels_CreateFcn(hObject, eventdata, handles)
% hObject    handle to fibrosis_pixels (see GCBO)
% eventdata  reserved - to be defined in a future version of MATLAB
% handles    empty - handles not created until after all CreateFcns called

% Hint: edit controls usually have a white background on Windows.
%       See ISPC and COMPUTER.
if ispc && isequal(get(hObject,'BackgroundColor'), get(0,'defaultUicontrolBackgroundColor'))
    set(hObject,'BackgroundColor','white');
end


function fibrosis_percent_Callback(hObject, eventdata, handles)
% hObject    handle to fibrosis_percent (see GCBO)
% eventdata  reserved - to be defined in a future version of MATLAB
% handles    structure with handles and user data (see GUIDATA)

% Hints: get(hObject,'String') returns contents of fibrosis_percent as text
%        str2double(get(hObject,'String')) returns contents of fibrosis_percent as a double


% --- Executes during object creation, after setting all properties.
function fibrosis_percent_CreateFcn(hObject, eventdata, handles)
% hObject    handle to fibrosis_percent (see GCBO)
% eventdata  reserved - to be defined in a future version of MATLAB
% handles    empty - handles not created until after all CreateFcns called

% Hint: edit controls usually have a white background on Windows.
%       See ISPC and COMPUTER.
if ispc && isequal(get(hObject,'BackgroundColor'), get(0,'defaultUicontrolBackgroundColor'))
    set(hObject,'BackgroundColor','white');
end


% --- Executes on button press in save_data.
function save_data_Callback(hObject, eventdata, handles)
% hObject    handle to save_data (see GCBO)
% eventdata  reserved - to be defined in a future version of MATLAB
% handles    structure with handles and user data (see GUIDATA)

flg_data_folder=handles.flg_data_folder;
if flg_data_folder==0
    msgbox("A folder for saving data is needed. Please select a folder for results");
end
if flg_data_folder==1
results_folder=handles.results_folder;
filname=handles.fname_ori;

csvoutput = [results_folder, '\fibrosis_results.csv'];
if exist(fullfile(results_folder, '\fibrosis_results.csv')) == 0
    title_str=cat(2,'File Name',',','Fibrosis pixels',',','Tissue Pixels',',','Fibrosis Percentage');
    fid=fopen(csvoutput,'at+');
    fprintf(fid,title_str);
    fprintf(fid,'\n');
    fclose(fid);

%     dlmwrite(csvoutput, title_str, 'delimiter', ' ', '-append');
end
out=cat(2,filname,',', get(handles.fibrosis_pixels,'String'),',',get(handles.tissue_pixels,'String'),',',get(handles.fibrosis_percent,'String'));
% dlmwrite(csvoutput, out, 'delimiter', ' ', '-append');
fid=fopen(csvoutput,'at+');
fprintf(fid,out);
fprintf(fid,'\n');
fclose(fid);
set(handles.save_data,'ForegroundColor','b');
end


function file_name_display_Callback(hObject, eventdata, handles)
% hObject    handle to file_name_display (see GCBO)
% eventdata  reserved - to be defined in a future version of MATLAB
% handles    structure with handles and user data (see GUIDATA)

% Hints: get(hObject,'String') returns contents of file_name_display as text
%        str2double(get(hObject,'String')) returns contents of file_name_display as a double


% --- Executes during object creation, after setting all properties.
function file_name_display_CreateFcn(hObject, eventdata, handles)
% hObject    handle to file_name_display (see GCBO)
% eventdata  reserved - to be defined in a future version of MATLAB
% handles    empty - handles not created until after all CreateFcns called

% Hint: edit controls usually have a white background on Windows.
%       See ISPC and COMPUTER.
if ispc && isequal(get(hObject,'BackgroundColor'), get(0,'defaultUicontrolBackgroundColor'))
    set(hObject,'BackgroundColor','white');
end


% --- Executes on button press in enlarge_original_image.
function enlarge_original_image_Callback(hObject, eventdata, handles)
% hObject    handle to enlarge_original_image (see GCBO)
% eventdata  reserved - to be defined in a future version of MATLAB
% handles    structure with handles and user data (see GUIDATA)

img_ori=handles.img_ori;
img_double=im2double(img_ori);
figure;
imshow(img_double)


% --- Executes on button press in pushbutton6.
function pushbutton6_Callback(hObject, eventdata, handles)
% hObject    handle to pushbutton6 (see GCBO)
% eventdata  reserved - to be defined in a future version of MATLAB
% handles    structure with handles and user data (see GUIDATA)

img_highlighted=handles.img_highlighted;

    figure;
    imshow(img_highlighted)
 


function tissue_thd_black_Callback(hObject, eventdata, handles)
% hObject    handle to tissue_thd_black (see GCBO)
% eventdata  reserved - to be defined in a future version of MATLAB
% handles    structure with handles and user data (see GUIDATA)

% Hints: get(hObject,'String') returns contents of tissue_thd_black as text
%        str2double(get(hObject,'String')) returns contents of tissue_thd_black as a double


% --- Executes during object creation, after setting all properties.
function tissue_thd_black_CreateFcn(hObject, eventdata, handles)
% hObject    handle to tissue_thd_black (see GCBO)
% eventdata  reserved - to be defined in a future version of MATLAB
% handles    empty - handles not created until after all CreateFcns called

% Hint: edit controls usually have a white background on Windows.
%       See ISPC and COMPUTER.
if ispc && isequal(get(hObject,'BackgroundColor'), get(0,'defaultUicontrolBackgroundColor'))
    set(hObject,'BackgroundColor','white');
end


function tissue_thd_white_Callback(hObject, eventdata, handles)
% hObject    handle to tissue_thd_white (see GCBO)
% eventdata  reserved - to be defined in a future version of MATLAB
% handles    structure with handles and user data (see GUIDATA)

% Hints: get(hObject,'String') returns contents of tissue_thd_white as text
%        str2double(get(hObject,'String')) returns contents of tissue_thd_white as a double


% --- Executes during object creation, after setting all properties.
function tissue_thd_white_CreateFcn(hObject, eventdata, handles)
% hObject    handle to tissue_thd_white (see GCBO)
% eventdata  reserved - to be defined in a future version of MATLAB
% handles    empty - handles not created until after all CreateFcns called

% Hint: edit controls usually have a white background on Windows.
%       See ISPC and COMPUTER.
if ispc && isequal(get(hObject,'BackgroundColor'), get(0,'defaultUicontrolBackgroundColor'))
    set(hObject,'BackgroundColor','white');
end


% --- Executes on button press in pushbutton7.
function pushbutton7_Callback(hObject, eventdata, handles)
% hObject    handle to pushbutton7 (see GCBO)
% eventdata  reserved - to be defined in a future version of MATLAB
% handles    structure with handles and user data (see GUIDATA)

img_tissue=handles.img_tissue;
figure;
imshow(img_tissue)


% --- Executes on button press in folder_results.
function folder_results_Callback(hObject, eventdata, handles)
% hObject    handle to folder_results (see GCBO)
% eventdata  reserved - to be defined in a future version of MATLAB
% handles    structure with handles and user data (see GUIDATA)

flg_data_folder=handles.flg_data_folder;
results_folder=uigetdir;

flg_data_folder=1;
handles.results_folder=results_folder;
handles.flg_data_folder=flg_data_folder;
guidata(hObject, handles);


% --- Executes on button press in bg_switch.
function bg_switch_Callback(hObject, eventdata, handles)
% hObject    handle to bg_switch (see GCBO)
% eventdata  reserved - to be defined in a future version of MATLAB
% handles    structure with handles and user data (see GUIDATA)

bg_indicator=handles.bg_indicator;
img_highlighted=handles.img_highlighted;
img_highlighted_white=img_highlighted;
row=handles.row_img;
col=handles.col_img;
if rem(bg_indicator,2)==0
    for b=1:1:row
      for c=1:1:col
         if img_highlighted_white(b,c,1)==0&&img_highlighted_white(b,c,2)==0&&img_highlighted_white(b,c,3)==0
             img_highlighted_white(b,c,1)=1;
             img_highlighted_white(b,c,2)=1;
             img_highlighted_white(b,c,3)=1;

         elseif img_highlighted_white(b,c,1)==0&&img_highlighted_white(b,c,2)==0&&img_highlighted_white(b,c,3)<0.2     
             img_highlighted_white(b,c,1)=1;
             img_highlighted_white(b,c,2)=1;
             img_highlighted_white(b,c,3)=1;
         end
      end
    end
    bg_indicator=bg_indicator+1;
   
    axes(handles.axes2)
    imshow(img_highlighted_white)
    title('Image With High Contrast')
    axis off 
handles.img_highlighted_white=img_highlighted_white;
elseif rem(bg_indicator,2)==1
    axes(handles.axes2)
    imshow(img_highlighted)
    title('Image With High Contrast')
    axis off 
    bg_indicator=bg_indicator+1;
end

handles.bg_indicator=bg_indicator;
handles.img_highlighted_white=img_highlighted_white;

 guidata(hObject, handles);
```
