## Supplemental codes for "Detection, Isolation and Quantification of Myocardial Infarct with Four Different Histological Staining Techniques": Instruction for fibrosis analysis.pdf

### The Instruction for Fibrosis Analysis in the MTS-Stained Tissue

#### Preparation

Two versions of this program are provided in the supplemental material. One version includes two program files in '.fig' and '.m' formats, while the other is an executable file in '.exe' format. To run either version of this program, you will need MATLAB software or MATLAB Runtime:

1. MATLAB software
   If MATLAB software is installed on your computer, you can directly open and run the files in '.fig' and '.m' formats using MATLAB.
2. MATLAB Runtime
   If the MATLAB software is not available on your computer, you can run the executable file in '.exe' format using MATLAB Runtime. Since this program is developed with MATLAB R2022b, the MATLAB Runtime version 9.13 is required. You can download it from this link for free: https://www.mathworks.com/products/compiler/matlab-runtime.html, or search 'MATLAB Runtime' on mathworks website.

#### Fibrosis analysis

1. Open and run *fibrosis_analyzer.m* in the MATLAB, or run *fibrosis_analyzer.exe* after MATLAB Runtime is installed (Figure 1).
2. Click 'Select a folder for results' to select a folder to save the data. This step can be done any time before clicking 'Save'.
3. Image loading. Click 'Load an image' to load an MTS-stained tissue section (Figure 2). This program supports images in '.tif', 'png', 'jpg' and 'gif' formats. The file name is displayed above the image. Click 'Zoom' to see the large version of the loaded image if needed.
4. Fibrosis extraction. 'Extraction Threshold' is a color-related threshold to isolate the fibrotic tissues (blue color) from the whole tissue section (Figure 3). The default threshold value is 1.2, but it can be changed. In this example, a value of 1.4 is used as it can greatly differentiate the fibrosis level in my tissues. Then, click 'Fibrosis Extraction' to extract all fibrotic tissues. The generated number of 'Fibrosis pixels' is shown in the right bottom. Click 'Zoom' to see the large version of the fibrosis image if needed.
5. Tissue extraction. 'Black Threshold' and 'White Threshold' are used to change the background color to black (Figure 4). Click 'Tissue Extraction' to show the tissue only in the black background. The generated number of 'Tissue pixels' is shown in the right bottom. Click 'Zoom' to see the large version of the tissue image if needed. When 'Fibrosis pixels' and 'Tissue pixels' are available, the percentage of the fibrosis relative to the tissue (Fibrosis(%)) can be determined (Figure 5).
6. Click 'Save' to save all these three data on the computer.

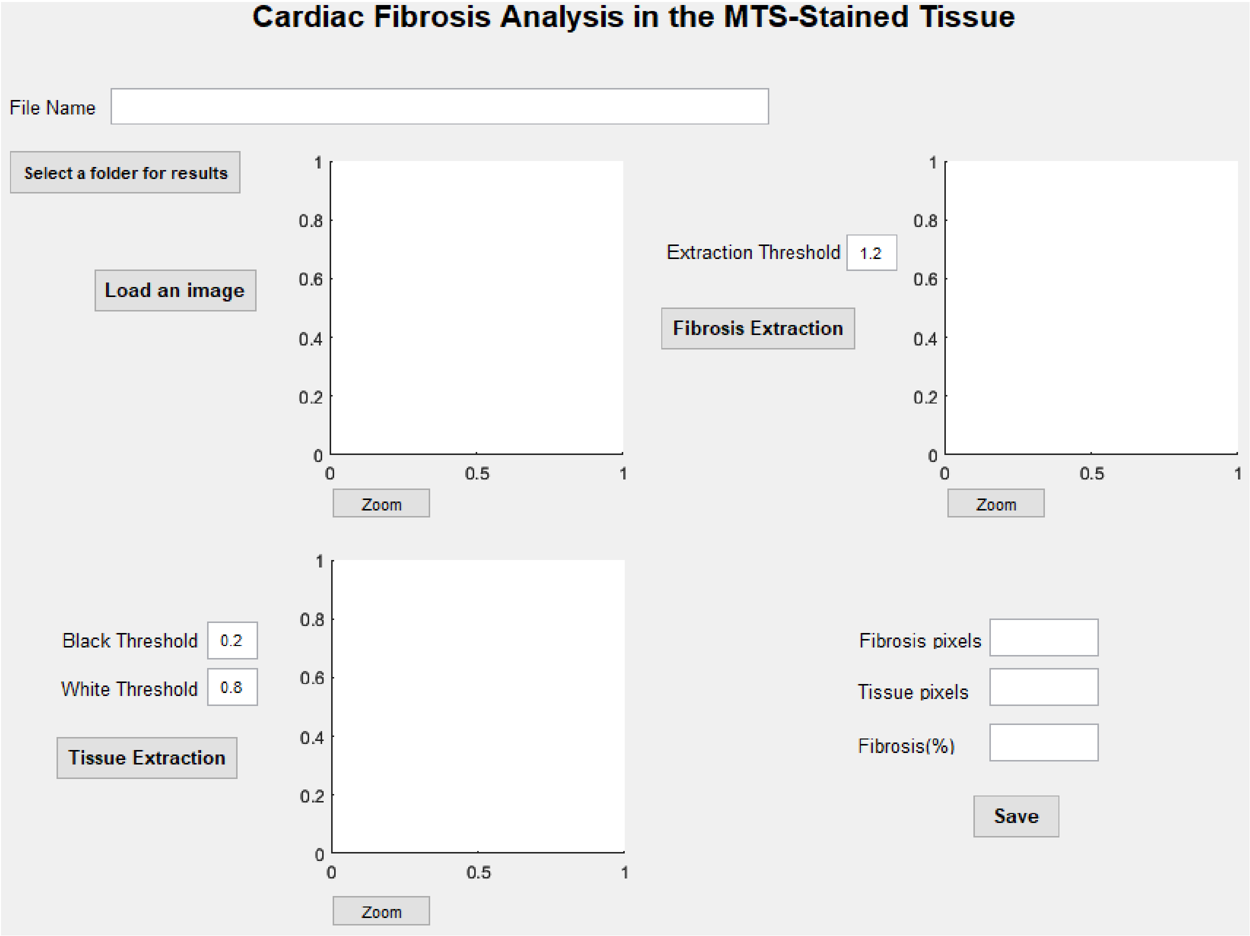


Figure 1

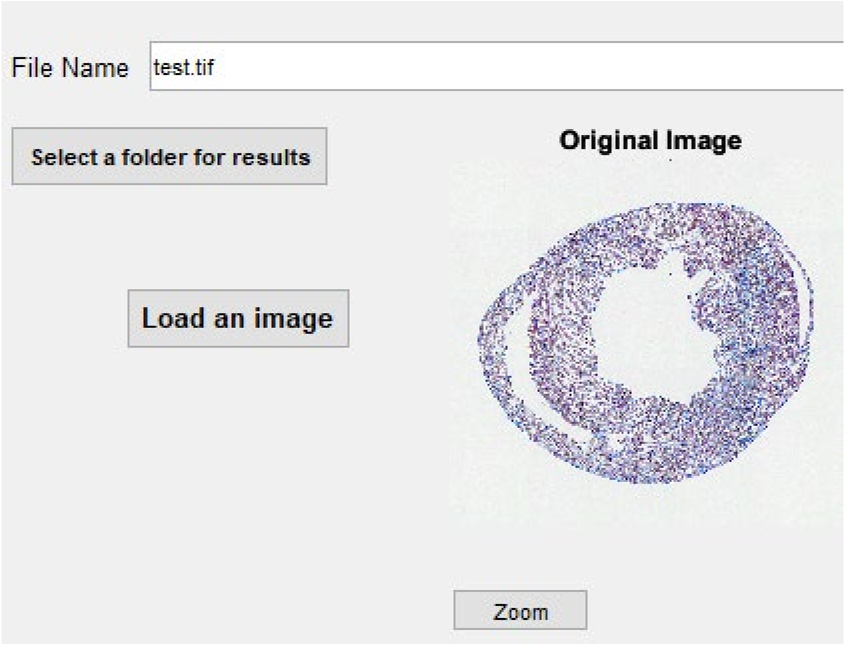


Figure 2

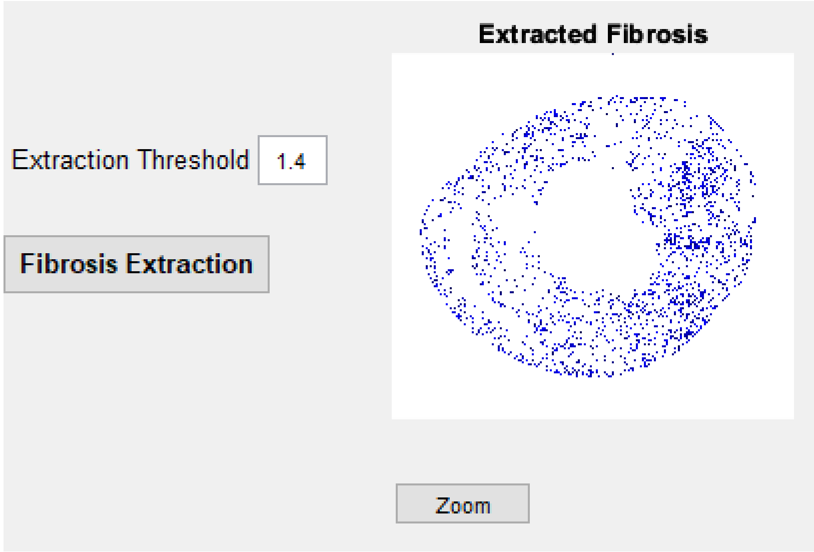


Figure 3

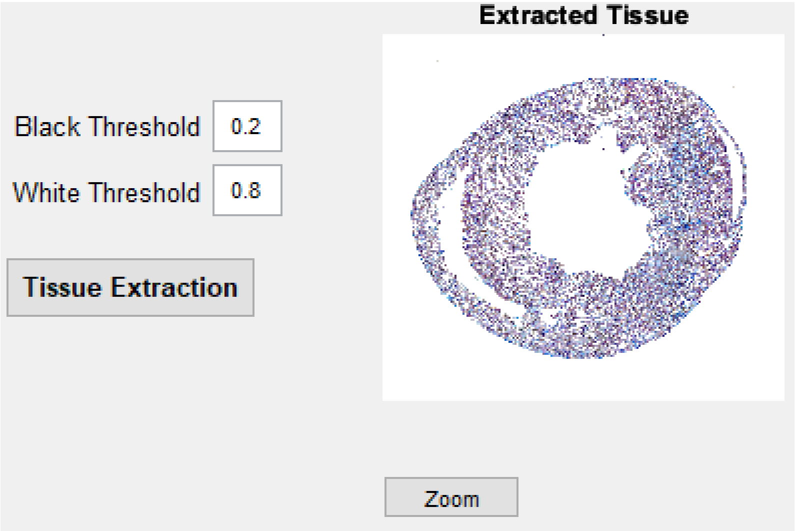


Figure 4

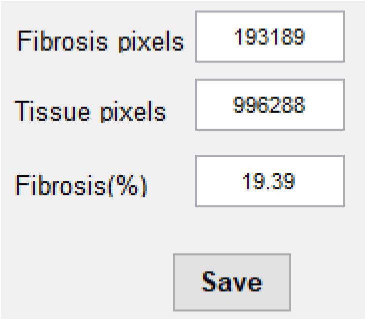


Figure 5
