## Supplemental codes for "Detection, Isolation and Quantification of Myocardial Infarct with Four Different Histological Staining Techniques": Instruction for image splitting.pdf

### The instruction for Image Splitting

#### Preparation

Two versions of this program are provided in the supplemental material. One version includes two program files in '.fig' and '.m' formats, while the other is an executable file in '.exe' format. To run either version of this program, you will need MATLAB software or MATLAB Runtime:

1. MATLAB software
   If MATLAB software is installed on your computer, you can directly open and run the files in '.fig' and '.m' formats using MATLAB.
2. MATLAB Runtime
   If the MATLAB software is not available on your computer, you can run the executable file in '.exe' format using MATLAB Runtime. Since this program is developed with MATLAB R2022b, the MATLAB Runtime version 9.13 is required. You can download it from this link for free: https://www.mathworks.com/products/compiler/matlab-runtime.html, or search 'MATLAB Runtime' on mathworks website.

#### Image splitting

1. Open and run *split_image.m* in the MATLAB, or run *split_image.exe* after MATLAB Runtime is installed (Figure 1).
2. Click 'Load an image' under the menu 'File'. This program supports any images in 'tif','jpg' or 'png' format. The loaded image can be seen on the left with the file name on the top (Figure 2).
3. Enter the amount of images to split and then click 'Confirm' (Figure 3). Then you will see lines with different colors on the original image (Figure 4). Those lines are used to determine the location of each image.
4. 'Top position' and 'Bottom position' are used to change the location of each line. Each pair of line with the same color determines the top and bottom sides of each image. Change those numbers and then click 'Confirm' (Figure 4) to see if the lines are placed properly.
5. Then click 'Split' to split all images. The split images can be seen on the right (Figure 5).
6. Then click 'Save as' to save all split images with the format 'tif' in the computer (Figure 5).

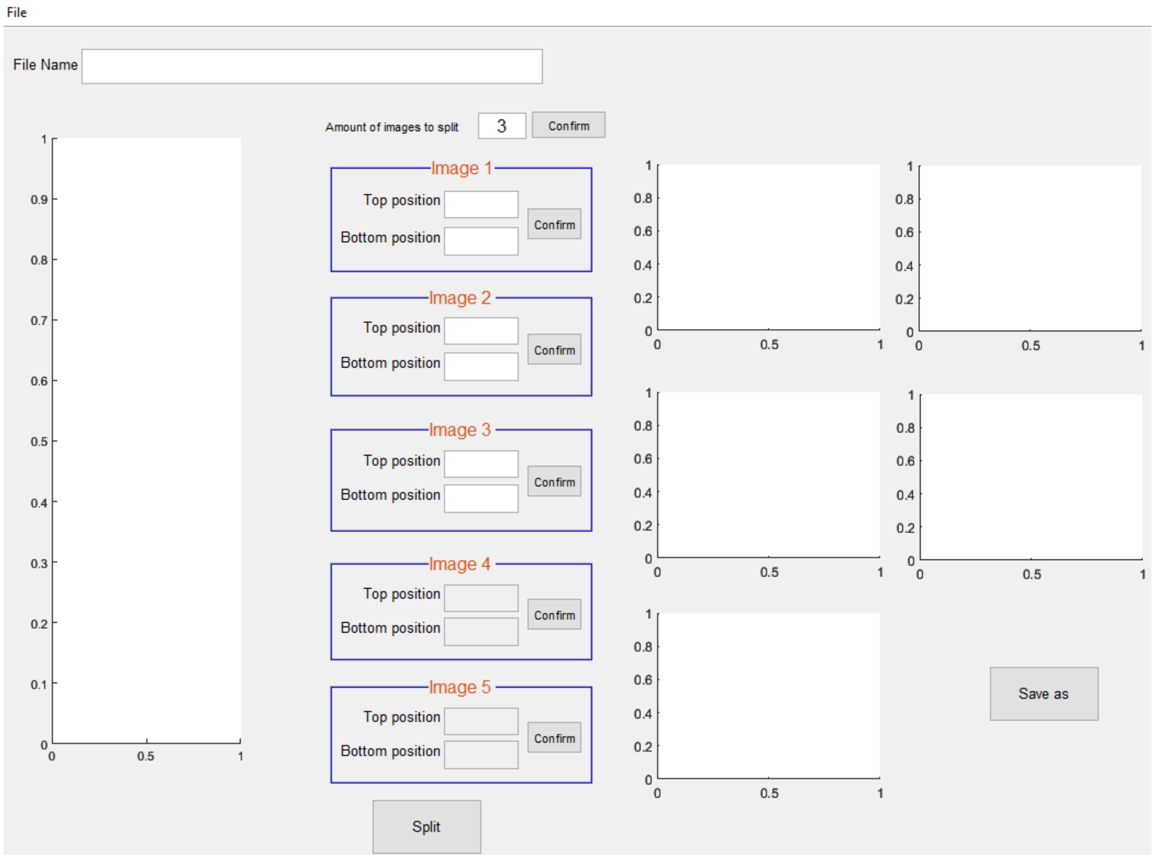
File
File Name
Amount of images to split
3
Confirm
Image 1
Top position
Bottom position
Confirm
Image 2
Top position
Bottom position
Confirm
Image 3
Top position
Bottom position
Confirm
Image 4
Top position
Bottom position
Confirm
Image 5
Top position
Bottom position
Confirm
Split
Save as

Figure 1

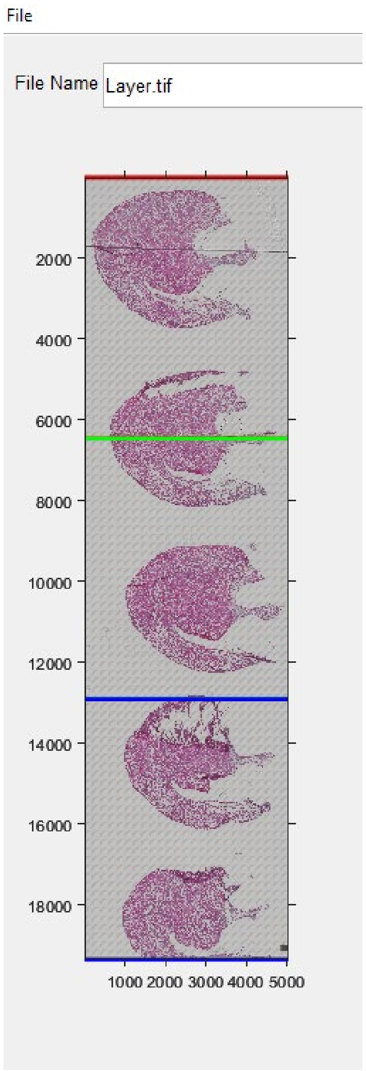


Figure 2

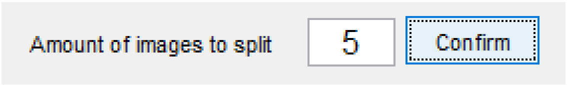


Figure 3

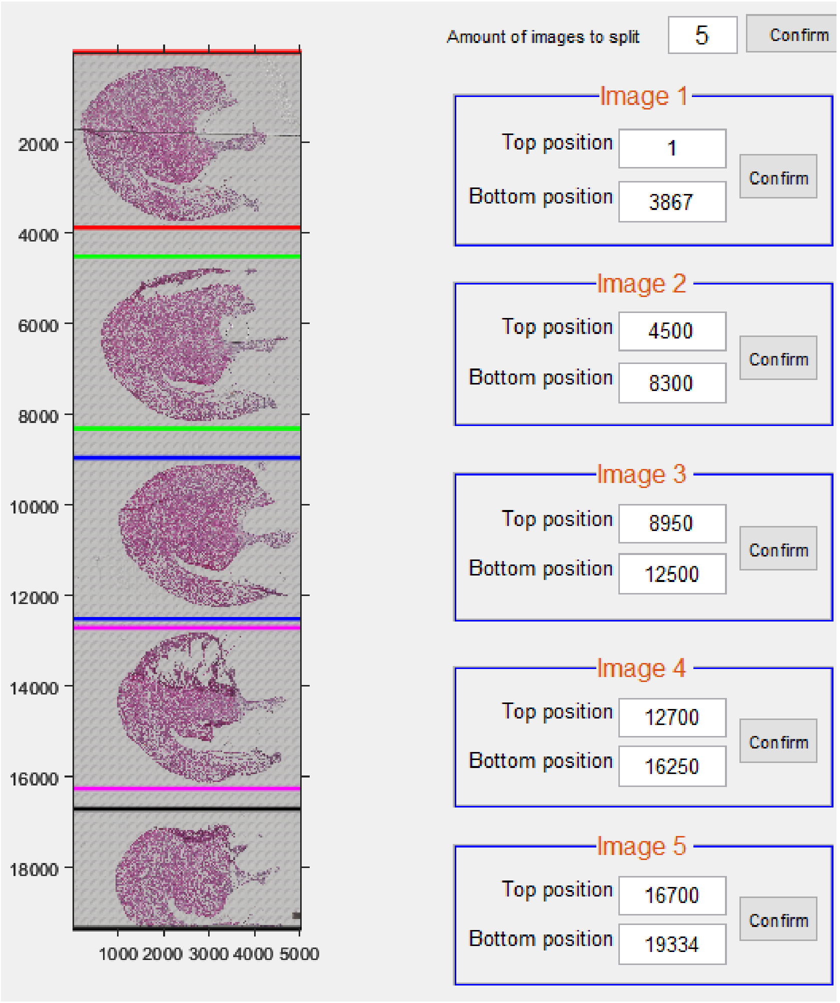
Amount of images to split
5
Confirm
Image 1
Top position
1
Bottom position
3867
Confirm
Image 2
Top position
4500
Bottom position
8300
Confirm
Image 3
Top position
8950
Bottom position
12500
Confirm
Image 4
Top position
12700
Bottom position
16250
Confirm
Image 5
Top position
16700
Bottom position
19334
Confirm
2000
4000
6000
8000
10000
12000
14000
16000
18000
1000 2000 3000 4000 5000

Figure 4

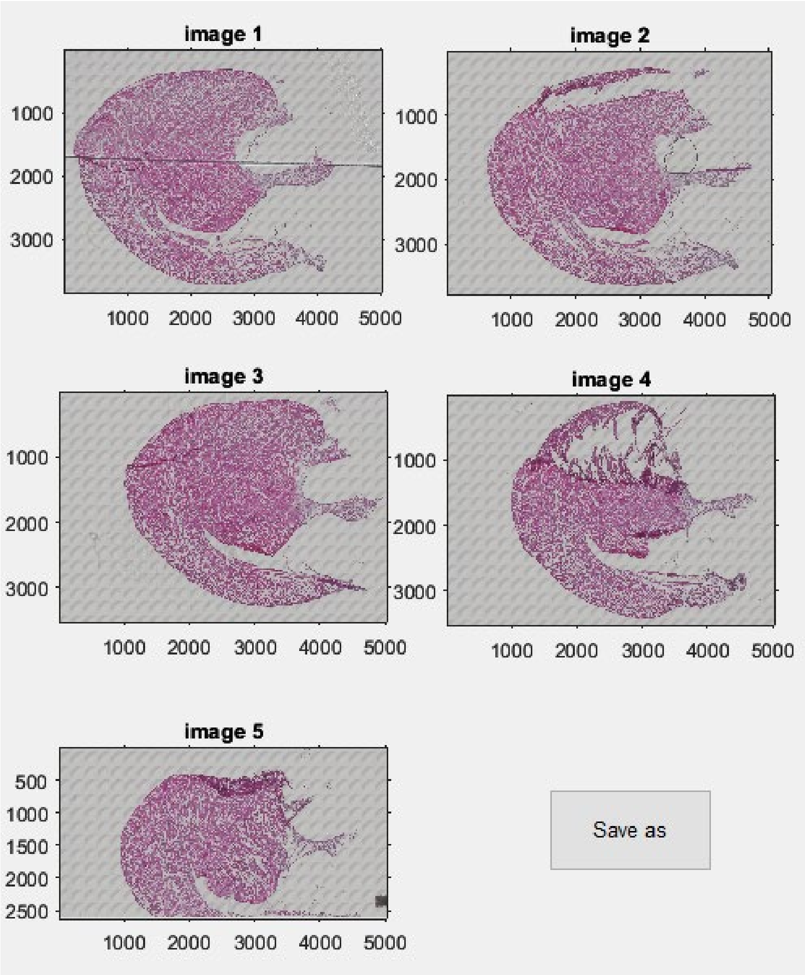

image 1
image 2
image 3
image 4
image 5
1000
2000
3000
4000
5000
500
1500
2500
Save as


Figure 5
