## Supplemental codes for "Detection, Isolation and Quantification of Myocardial Infarct with Four Different Histological Staining Techniques": Instruction for scar size analysis.pdf

### The Instruction for Scar Tissue Size Analyzer

#### Preparation

Two versions of this program are provided in the supplemental material. One version includes two program files in '.fig' and '.m' formats, while the other is an executable file in '.exe' format. To run either version of this program, you will need MATLAB software or MATLAB Runtime:

1. MATLAB software
   If MATLAB software is installed on your computer, you can directly open and run the files in '.fig' and '.m' formats using MATLAB.
2. MATLAB Runtime
   If the MATLAB software is not available on your computer, you can run the executable file in '.exe' format using MATLAB Runtime. Since this program is developed with MATLAB R2022b, the MATLAB Runtime version 9.13 is required. You can download it from this link for free: https://www.mathworks.com/products/compiler/matlab-runtime.html, or search 'MATLAB Runtime' on mathworks website.

#### Scar analysis

1. Open and run *scar_size_analysis.m* in the MATLAB, or run *scar_size_analysis.exe* after MATLAB Runtime is installed (Figure 1).
2. On the interface of this program, you will find four different staining methods on the top left with MTS (default), H&E, TTC and PSR. Select the appropriate one for your image. Here we use MTS as an example to show how to use this program to analyze the myocardial infarct or scar.
3. Click 'Load' to load an image in '.tif', 'png', 'jpg' or 'gif' format. The file name is shown above the image (Figure 2). The pixel size can be entered right now. If the pixel size is not needed, leave '1' in the box. Click 'Zoom' to see the large version of the loaded image if needed.
4. The 'Extraction_thd' is the color-related threshold used to identify and isolate the scar (Figure 3). The pre-determined value varies with the staining method and it can be changed to get the good identification of the scar. Click 'Highlight the scar' to show the identified scar image. The default background is black. Click 'Background' to toggle the black and white background if needed. Click 'Zoom' to see the large version of the scar image if needed. Click 'Save highlighted image' to save the scar image. Before you click this button, you may need to select a folder to save the image by clicking 'Select a folder for saving images' (Figure 4). The folder can be selected any time before saving any images.
5. Click 'Extract the Scar' to get the final scar image before masking (Figure 5). Click 'Save extracted image' to save the image if needed.
6. Click 'Select ROI' to show the large version of the image produced in the last step. When the mouse cursor is in the image, it will be changed to '+'. Then press and

hold the left click and drag from left top to right bottom to select the scar location (Figure 6). Right click in the selected scar range and select ‘Crop Image’.

7. The cropped scar image will be shown with a title ‘Selected Scar’ (Figure 7). Click ‘Zoom’ to see the large version of the cropped scar image. Click ‘Save selected ROI’ to save the image if needed.
8. Mask the scar (Figure 8). Three different thresholds are used to mask the scar. ‘Size_thd’ and ‘Extract_thd’ are used to remove the faint blue colors to make sure only the scar is isolated for masking. ‘Dilation_thd’ is used to dilate and mask the scar region. The values for these three thresholds vary with the staining method. Click ‘Mask’ to show the masked scar. Click ‘Overlap’ to check the masking quality by overlapping the scar image and the masked image. Click ‘Zoom’ to see the large version of the masked scar image. Click ‘Save masked scar’ to save the image if needed.
9. After clicking ‘Mask’, the number of ‘Scar Pixels’ is displayed in the panel of ‘Scar info’ (Figure 9). If the pixel size is available, the calculated ‘Scar Area (mm^2)’ is also displayed. If the pixel size is ‘1’, the ‘Scar Area (mm^2)’ will be the same as the ‘Scar Pixels’. If the produced scar pixels and scar area will be saved at this step, make sure to click ‘Select a folder for saving results’ to select a folder for saving the data (Figure 4).
10. Mask the whole tissue section (Figure 10). Up to five thresholds are used to mask the whole tissue section. These threshold values vary with the staining method. ‘Remove_BG1’ and ‘Remove_BG2’ are used to set the background as black color. ‘Size_thd1’ and ‘Size_thd2’ are used to remove small pieces not belonging to the tissue. ‘Dilation_thd’ is used to dilate and mask the tissue section. Click ‘Mask’ to mask the whole tissue section. Click ‘Zoom’ to see the large version of the masked whole tissue section image. Click ‘Save masked image’ to save the image.
11. After clicking ‘Mask’, the number of ‘Tissue Pixels’ is displayed in the panel of ‘Scar info’ (Figure 11). If the pixel size is available, the calculated Tissue Area (mm^2)’ is also displayed. If the pixel size is ‘1’, the ‘Tissue Area (mm^2)’ will be the same as the ‘Tissue Pixels’. If the produced Tissue pixels and Tissue area will be saved at this step, make sure to click ‘Select a folder for saving results’ to select a folder for saving the data (Figure 4).
12. When the ‘Scar Pixels’ and ‘Tissue Pixels’ are available, the percentage of the scar relative to the whole tissue section (Scar(%)) will be calculated and displayed in the panel of ‘Scar info’.
13. Click ‘Save’ in the panel of ‘Scar info’ to save all data in the computer.

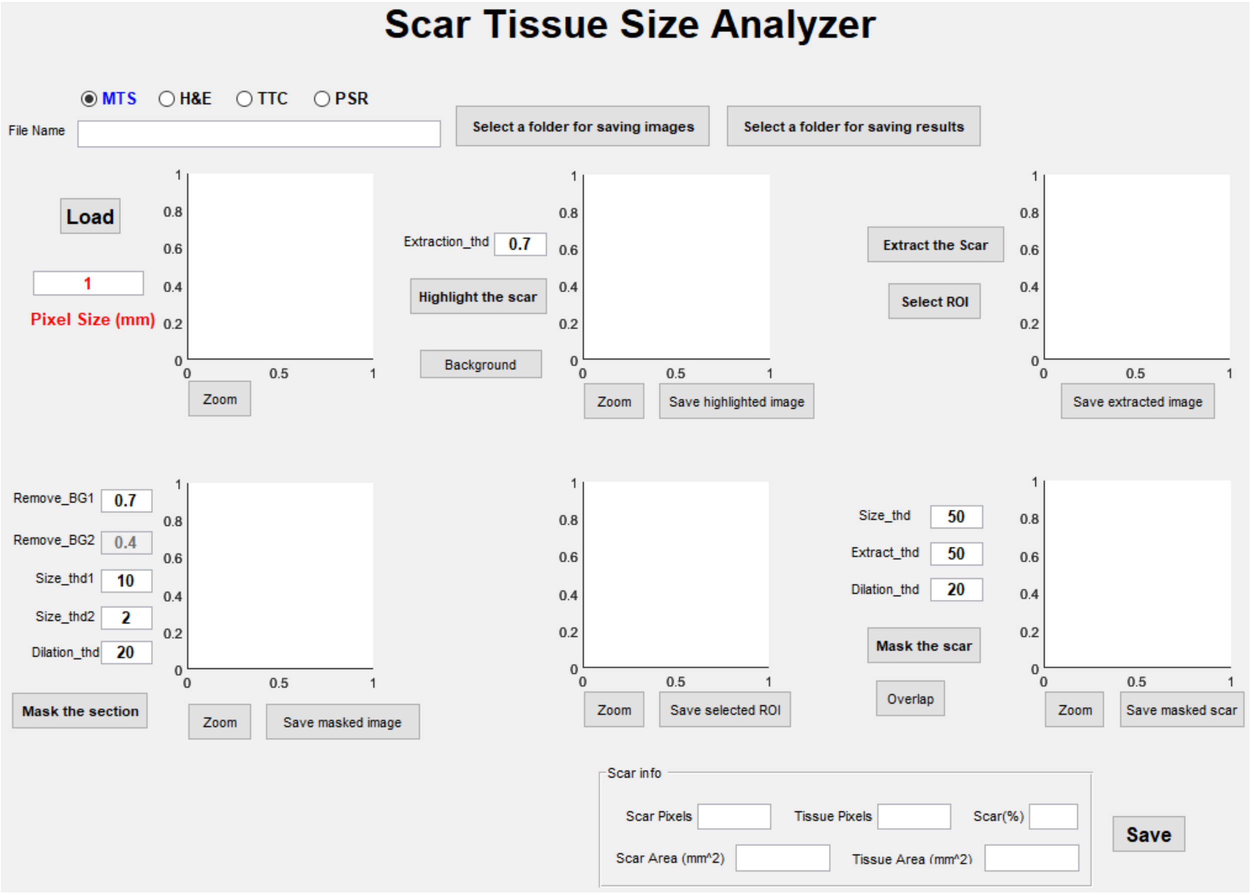


Figure 1

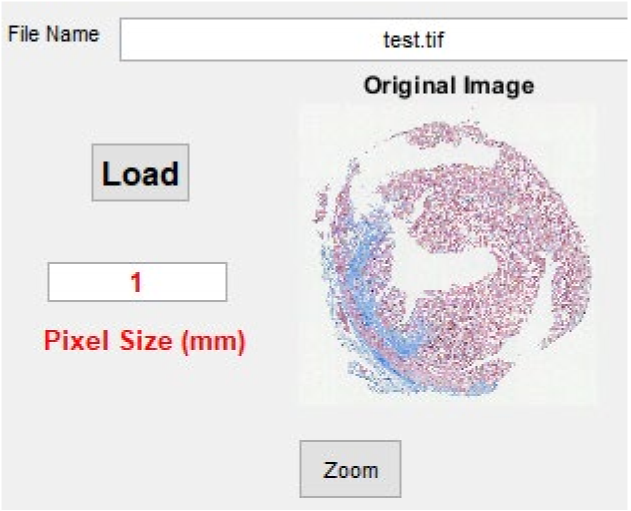


Figure 2

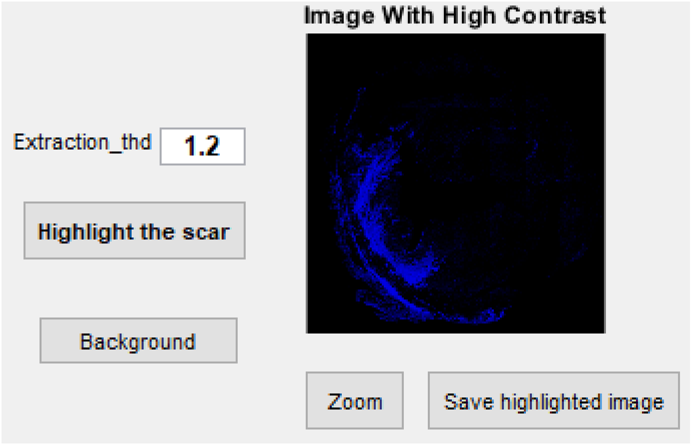


Figure 3

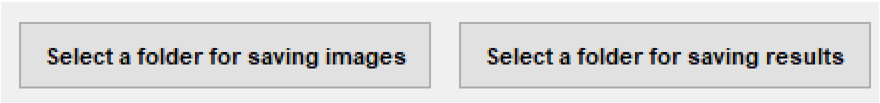


Figure 4

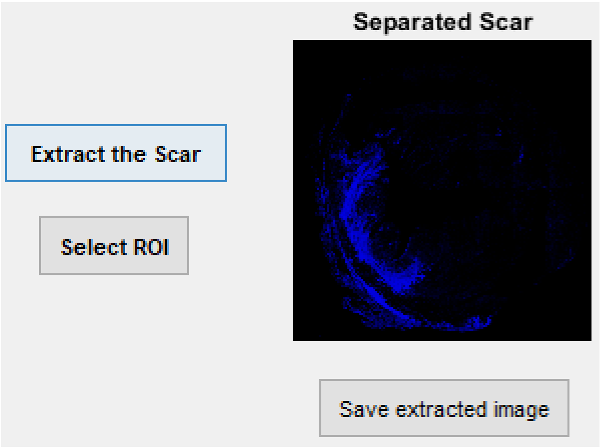


Figure 5

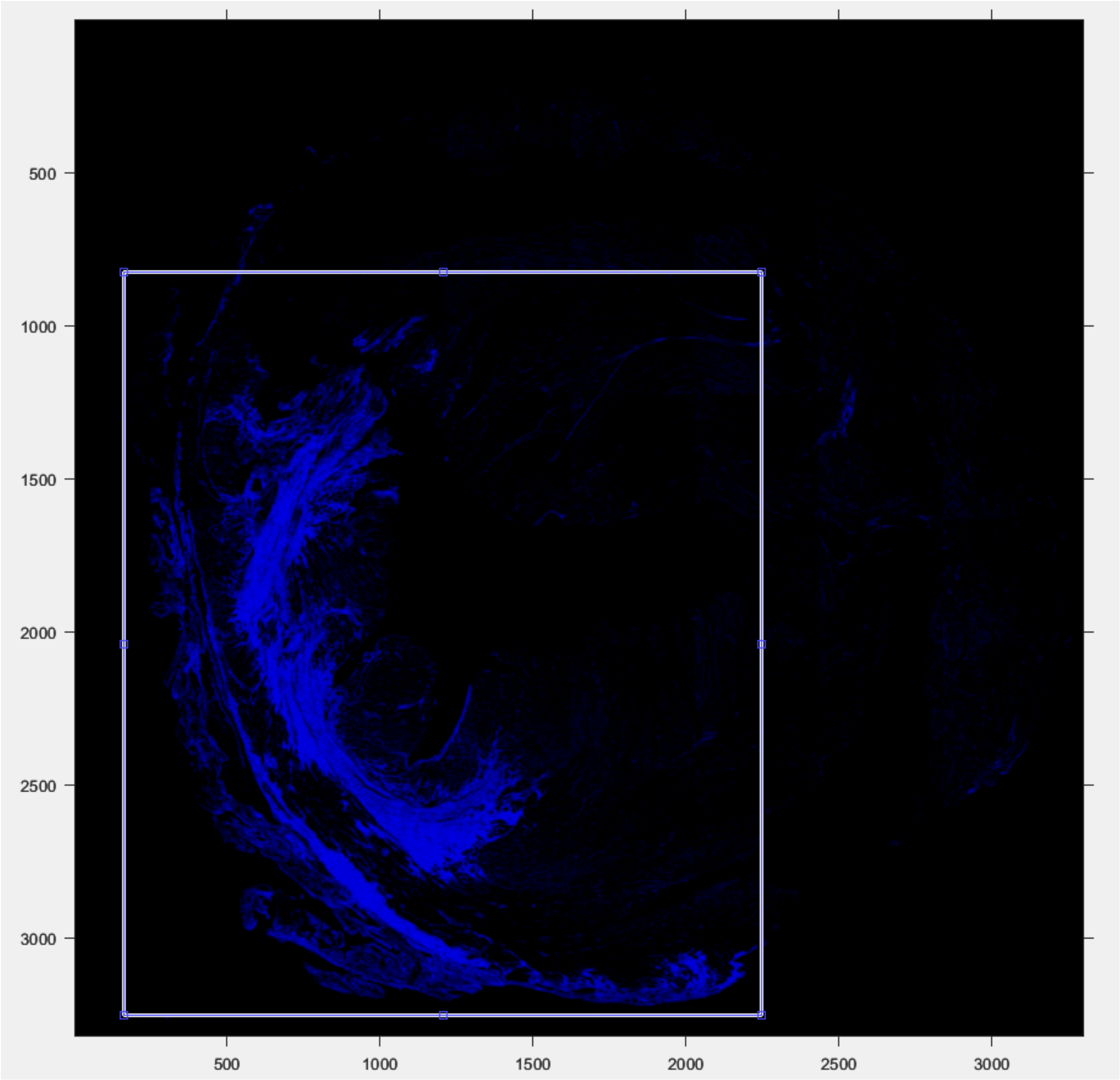


Figure 6

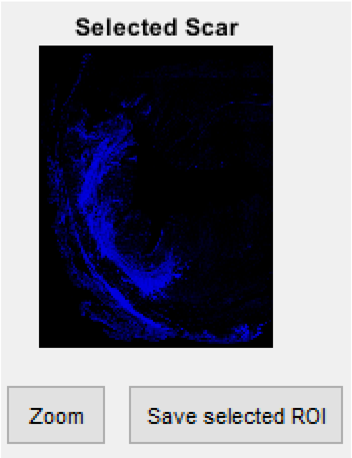


Figure 7

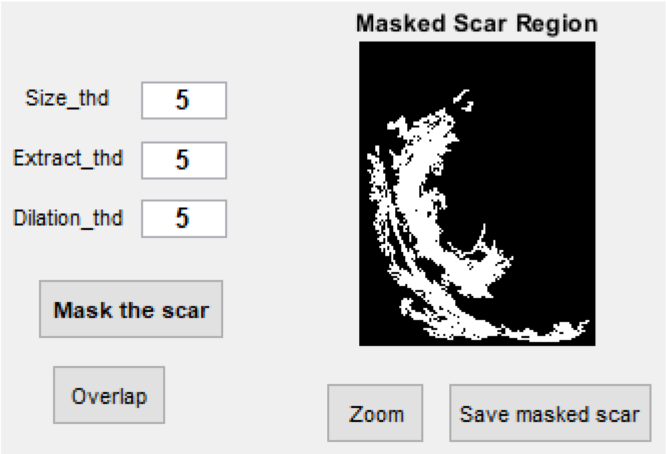


Figure 8

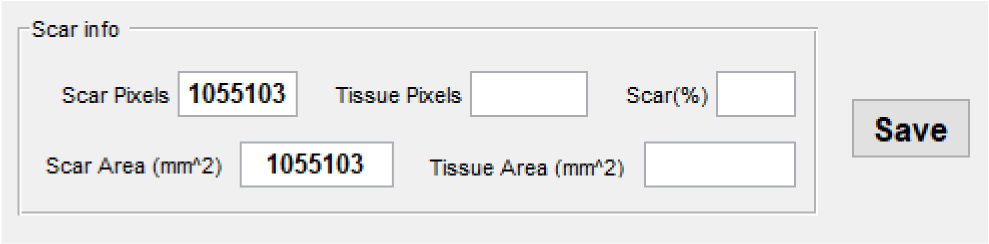


Figure 9

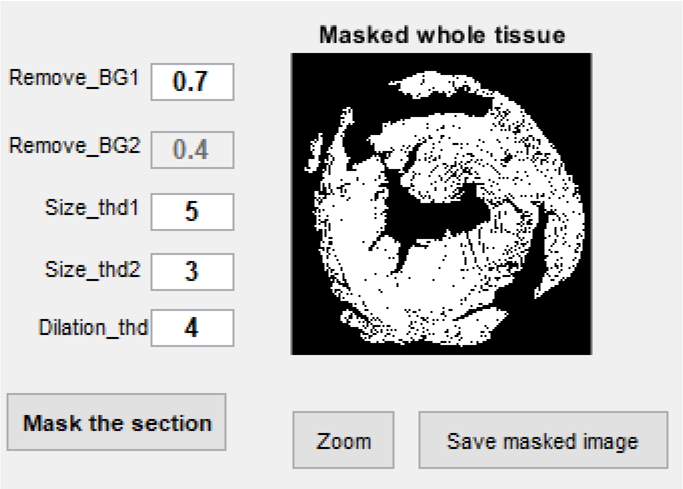


Figure 10

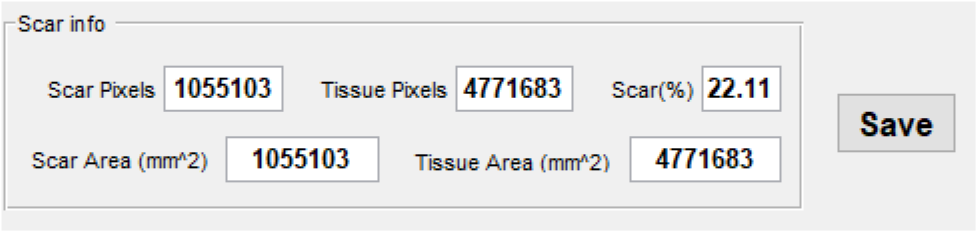


Figure 11
