## Supplemental codes for "Detection, Isolation and Quantification of Myocardial Infarct with Four Different Histological Staining Techniques": scar_size_analysis.m

```
function varargout = scar_size_analysis(varargin)
% This script was completed by Xiaobo Wu with the suggestions from Tom Liu
% and Linnea Meier
% SCAR_SIZE_ANALYSIS MATLAB code for scar_size_analysis.fig
%      SCAR_SIZE_ANALYSIS, by itself, creates a new SCAR_SIZE_ANALYSIS or raises the existing
%      singleton*.
%
%      H = SCAR_SIZE_ANALYSIS returns the handle to a new SCAR_SIZE_ANALYSIS or the handle to
%      the existing singleton*.
%
%      SCAR_SIZE_ANALYSIS('CALLBACK',hObject,eventData,handles,...) calls the local
%      function named CALLBACK in SCAR_SIZE_ANALYSIS.M with the given input arguments.
%
%      SCAR_SIZE_ANALYSIS('Property','Value',...) creates a new SCAR_SIZE_ANALYSIS or raises the
%      existing singleton*.  Starting from the left, property value pairs are
%      applied to the GUI before scar_size_analysis_OpeningFcn gets called.  An
%      unrecognized property name or invalid value makes property application
%      stop.  All inputs are passed to scar_size_analysis_OpeningFcn via varargin.
%
%      *See GUI Options on GUIDE's Tools menu.  Choose "GUI allows only one
%      instance to run (singleton)".
%
% See also: GUIDE, GUIDATA, GUIHANDLES

% Edit the above text to modify the response to help scar_size_analysis

% Last Modified by GUIDE v2.5 15-Jul-2024 12:25:40

% Begin initialization code - DO NOT EDIT
gui_Singleton = 1;
gui_State = struct('gui_Name',       mfilename, ...
                   'gui_Singleton',  gui_Singleton, ...
                   'gui_OpeningFcn', @scar_size_analysis_OpeningFcn, ...
                   'gui_OutputFcn',  @scar_size_analysis_OutputFcn, ...
                   'gui_LayoutFcn',  [] , ...
                   'gui_Callback',   []);
if nargin && ischar(varargin{1})
    gui_State.gui_Callback = str2func(varargin{1});
end

if nargout
    [varargout{1:nargout}] = gui_mainfcn(gui_State, varargin{:});
else
    gui_mainfcn(gui_State, varargin{:});
end
% End initialization code - DO NOT EDIT


% --- Executes just before scar_size_analysis is made visible.
function scar_size_analysis_OpeningFcn(hObject, eventdata, handles, varargin)
% This function has no output args, see OutputFcn.
% hObject    handle to figure
% eventdata  reserved - to be defined in a future version of MATLAB
% handles    structure with handles and user data (see GUIDATA)
% varargin   command line arguments to scar_size_analysis (see VARARGIN)

% Choose default command line output for scar_size_analysis

% javaFrame = get(hObject,'JavaFrame');
% javaFrame.setFigureIcon(javax.swing.ImageIcon('icon_scar.png'));


flg_scar_folder=0;
flg_data_folder=0;
bg_indicator=0;
set(handles.mts_staining,'Value',1);
set(handles.mts_staining,'ForegroundColor','b');
set(handles.he_staining,'Value',0);
set(handles.ttc_staining,'Value',0);
set(handles.psr_staining,'Value',0);
% set(handles.axes1_original,'YTick',[],'XTick',[]);
% set(handles.axes2_highlighted,'YTick',[],'XTick',[]);
% set(handles.axes2_highlighted2,'YTick',[],'XTick',[]);
% set(handles.axes3_scar,'YTick',[],'XTick',[]);
% set(handles.axes4_dilated_scar,'YTick',[],'XTick',[]);
set(handles.bg_thd_whole,'String', '0.7');
set(handles.size_whole_thd1, 'String','10');
set(handles.size_whole_thd2, 'String','2');
set(handles.dilation_value_whole, 'String','20');


set(handles.bg_thd_ttc,'enable','off')

handles.flg_scar_folder=flg_scar_folder;
handles.flg_data_folder=flg_data_folder;
handles.output = hObject;
handles.bg_indicator = bg_indicator;
% Update handles structure
guidata(hObject, handles);

% UIWAIT makes scar_size_analysis wait for user response (see UIRESUME)
% uiwait(handles.figure1);


% --- Outputs from this function are returned to the command line.
function varargout = scar_size_analysis_OutputFcn(hObject, eventdata, handles) 
% varargout  cell array for returning output args (see VARARGOUT);
% hObject    handle to figure
% eventdata  reserved - to be defined in a future version of MATLAB
% handles    structure with handles and user data (see GUIDATA)

% Get default command line output from handles structure
varargout{1} = handles.output;


% --- Executes on button press in enlarge_original.
function enlarge_original_Callback(hObject, eventdata, handles)
% hObject    handle to enlarge_original (see GCBO)
% eventdata  reserved - to be defined in a future version of MATLAB
% handles    structure with handles and user data (see GUIDATA)
img_ori=handles.img_ori;
img_double=im2double(img_ori);
figure;
imshow(img_double)

% --- Executes on button press in enlarge_highlighted.
function enlarge_highlighted_Callback(hObject, eventdata, handles)
% hObject    handle to enlarge_highlighted (see GCBO)
% eventdata  reserved - to be defined in a future version of MATLAB
% handles    structure with handles and user data (see GUIDATA)
bg_indicator=handles.bg_indicator;
if rem(bg_indicator,2)==0

img_highlighted=handles.img_highlighted;
figure;
imshow(img_highlighted)
elseif rem(bg_indicator,2)==1
    img_highlighted_white=handles.img_highlighted_white;
    figure;
imshow(img_highlighted_white)
end

% --- Executes on button press in enlarge_scar.
function enlarge_scar_Callback(hObject, eventdata, handles)
% hObject    handle to enlarge_scar (see GCBO)
% eventdata  reserved - to be defined in a future version of MATLAB
% handles    structure with handles and user data (see GUIDATA)
scar_cropped=handles.scar_cropped;
figure;
imshow(scar_cropped)

% --- Executes on button press in enlarge_dilated_scar.
function enlarge_dilated_scar_Callback(hObject, eventdata, handles)
% hObject    handle to enlarge_dilated_scar (see GCBO)
% eventdata  reserved - to be defined in a future version of MATLAB
% handles    structure with handles and user data (see GUIDATA)
scar_dilated=handles.scar_dilated;
figure;
imshow(scar_dilated)


function color_thd_Callback(hObject, eventdata, handles)
% hObject    handle to color_thd (see GCBO)
% eventdata  reserved - to be defined in a future version of MATLAB
% handles    structure with handles and user data (see GUIDATA)

% Hints: get(hObject,'String') returns contents of color_thd as text
%        str2double(get(hObject,'String')) returns contents of color_thd as a double


% --- Executes during object creation, after setting all properties.
function color_thd_CreateFcn(hObject, eventdata, handles)
% hObject    handle to color_thd (see GCBO)
% eventdata  reserved - to be defined in a future version of MATLAB
% handles    empty - handles not created until after all CreateFcns called

% Hint: edit controls usually have a white background on Windows.
%       See ISPC and COMPUTER.
if ispc && isequal(get(hObject,'BackgroundColor'), get(0,'defaultUicontrolBackgroundColor'))
    set(hObject,'BackgroundColor','white');
end


% --- Executes on button press in scar_hl1.
function scar_hl1_Callback(hObject, eventdata, handles)
% hObject    handle to scar_hl1 (see GCBO)
% eventdata  reserved - to be defined in a future version of MATLAB
% handles    structure with handles and user data (see GUIDATA)
%-----pixels change from integer to double
img_ori=handles.img_ori;
img_double=im2double(img_ori);

%-----threadhold is got
con_thd=str2num(get(handles.color_thd,'string'));

[row,col,clr]=size(img_double);
img_thd=zeros(row,col,clr);
if get(handles.he_staining,'Value')==1

for a=1:1:clr
    for b=1:1:row
      for c=1:1:col
         if img_double(b,c,a)>con_thd
            img_thd(b,c,a)=0;
         elseif img_double(b,c,a)<=con_thd
             img_thd(b,c,a)=img_double(b,c,a);
         end
      end
    end
end


%------------filter-----------------

h=fspecial('disk',5);
img_filtered=imfilter(img_thd,h);
end

if get(handles.mts_staining,'Value')==1

    for b=1:1:row
        for c=1:1:col

           if img_double(b,c,3)/img_double(b,c,1)>=con_thd
               img_thd(b,c,1)=0;
               img_thd(b,c,2)=0;
               img_thd(b,c,3)=img_double(b,c,3);
           end
           if img_double(b,c,3)/img_double(b,c,1)<con_thd
               img_thd(b,c,1)=0;
               img_thd(b,c,2)=0;
               img_thd(b,c,3)=0;
           end
        end
         
    end

%     for b=1:1:row
%         for c=1:1:col
%           
%             if img_thd(b,c,3)>0.7
%             img_thd(b,c,3)=1;
%             end
%         end
%          
%     end

h=fspecial('disk',5);
img_filtered=imfilter(img_thd,h);
 

end

if get(handles.ttc_staining,'Value')==1
 
    %------remove background
for b=1:1:row
        for c=1:1:col

        if img_double(b,c,1)>0.75||img_double(b,c,2)>0.75||img_double(b,c,3)>0.75
               img_thd(b,c,1)=0;
               img_thd(b,c,2)=0;
               img_thd(b,c,3)=0;
        else
            img_thd(b,c,1)=img_double(b,c,1);
            img_thd(b,c,2)=img_double(b,c,2);    
            img_thd(b,c,3)=img_double(b,c,3);
        end
        end  
end
%----remove weak signals
 for b=1:1:row
        for c=1:1:col

        if img_thd(b,c,1)<0.3&&img_thd(b,c,2)<0.3&&img_thd(b,c,3)<0.3
               img_thd(b,c,1)=0;
               img_thd(b,c,2)=0;
               img_thd(b,c,3)=0;
        else
            img_thd(b,c,1)=img_thd(b,c,1);
            img_thd(b,c,2)=img_thd(b,c,2);    
            img_thd(b,c,3)=img_thd(b,c,3);
        end
        end  
 end

 for b=1:1:row
        for c=1:1:col

        if max(img_thd(b,c,:))<0.5
               img_thd(b,c,1)=0;
               img_thd(b,c,2)=0;
               img_thd(b,c,3)=0;
        
        end
        end  
 end

 
%----remove RGB color with large differences


std_thd=str2num(get(handles.color_thd,'String'));
for b=1:1:row
        for c=1:1:col
            if std(img_thd(b,c,:))>std_thd
               img_thd(b,c,1)=0;
               img_thd(b,c,2)=0;
               img_thd(b,c,3)=0;
            end
        end  
end


    %------R is larger than G or B in scar region
    
    for b=1:1:row
        for c=1:1:col


        if img_thd(b,c,2)>img_thd(b,c,1)&&img_thd(b,c,3)>img_thd(b,c,1)
               img_thd(b,c,1)=0;
               img_thd(b,c,2)=0;
               img_thd(b,c,3)=0;
        end
        end  
    end
    
    % h=fspecial('disk',5);
% img_filtered=imfilter(img_thd,h);
  img_filtered=img_thd;  
    
  
  
  
end

if get(handles.psr_staining,'Value')==1
    
    

%-----------change background to black
for b=1:1:row
        for c=1:1:col

        if img_double(b,c,2)>0.8||img_double(b,c,3)>0.8
               img_thd(b,c,1)=0;
               img_thd(b,c,2)=0;
               img_thd(b,c,3)=0;
               
        elseif img_double(b,c,1)<0.5
            
            img_thd(b,c,1)=0;
            img_thd(b,c,2)=0;    
            img_thd(b,c,3)=0;
        else
            img_thd(b,c,1)=img_double(b,c,1);
            img_thd(b,c,2)=img_double(b,c,2);
            img_thd(b,c,3)=img_double(b,c,3);
        end
        end  
end

%--------extract scar
for b=1:1:row
        for c=1:1:col
            temp=(img_thd(b,c,2)+img_thd(b,c,3))/2;
        if img_thd(b,c,1)-temp<0||img_thd(b,c,1)-temp<con_thd
            img_thd(b,c,1)=0;
            img_thd(b,c,2)=0;    
            img_thd(b,c,3)=0;
        end

        end
end


img_filtered=img_thd;  

end

axes(handles.axes2_highlighted)
imshow(img_filtered)
title('Image With High Contrast')
axis off
handles.img_highlighted=img_filtered;
handles.row_img=row;
handles.col_img=col;
handles.bg_indicator=0;
guidata(hObject, handles);


% --- Executes on button press in img_crop.
function img_crop_Callback(hObject, eventdata, handles)
% hObject    handle to img_crop (see GCBO)
% eventdata  reserved - to be defined in a future version of MATLAB
% handles    structure with handles and user data (see GUIDATA)
img_scar=handles.img_scar;

figure;
imshow(img_scar)
scar_cropped = imcrop(img_scar);
close gcf
axes(handles.axes3_scar)
imshow(scar_cropped)
title("Selected Scar")
axis off
handles.scar_cropped=scar_cropped;
guidata(hObject, handles);

% --- Executes on button press in scar_mask.
function scar_mask_Callback(hObject, eventdata, handles)
% hObject    handle to scar_mask (see GCBO)
% eventdata  reserved - to be defined in a future version of MATLAB
% handles    structure with handles and user data (see GUIDATA)
 scar_cropped=handles.scar_cropped;
% scar_cropped=imread('scar_cropped.tif');
grayImage=scar_cropped;
[rows, columns, numberOfColorChannels] = size(grayImage);
if numberOfColorChannels > 1
  grayImage = 255*rgb2gray(grayImage);
end
% 

if get(handles.he_staining,'Value')==1
mask_thd1=str2num(get(handles.thd1,'string'));
  mask = grayImage>mask_thd1;
% % 
 
mask_1 = bwareaopen(mask, 200);
% % % Get rid of white frame around outside border.
%   mask_2 = imclearborder(mask_1);
% % 
mask_thd2=str2num(get(handles.thd2,'string'));


  newmask = bwareafilt(mask_1, mask_thd2);
  img_filled_2 = imfill(newmask,8,'holes');
%    se = strel('line',30,80);
dilation_factor=str2num(get(handles.thd3,'string'));
   se = strel('square',dilation_factor);
  img_dilation = imdilate(newmask,se);
  scar_dilated = imfill(img_dilation,8,'holes');
end

if get(handles.mts_staining,'Value')==1
   mask_thd1=str2num(get(handles.thd1,'string'));
   mask_thd2=str2num(get(handles.thd2,'string'));
   mask = grayImage>mask_thd1;
    mask1 = bwareafilt(mask, mask_thd2);
    dilation_factor=str2num(get(handles.thd3,'string'));
   mask = bwareaopen(grayImage,mask_thd1);
    se = strel('square',dilation_factor);
  scar_dilated = imdilate(mask1,se);
 
end
   
   if get(handles.ttc_staining,'Value')==1
   mask_thd1=str2num(get(handles.thd1,'string'));
   mask_thd2=str2num(get(handles.thd2,'string'));
   mask = bwareaopen(grayImage,mask_thd1);
    mask1 = bwareafilt(mask, mask_thd2);
    dilation_factor=str2num(get(handles.thd3,'string'));
   mask = bwareaopen(grayImage,mask_thd1);
    se = strel('square',dilation_factor);
  scar_dilated = imdilate(mask1,se);
 
   end

 if get(handles.psr_staining,'Value')==1
   mask_thd1=str2num(get(handles.thd1,'string'));
   mask_thd2=str2num(get(handles.thd2,'string'));
   mask = bwareaopen(grayImage,mask_thd1);
    mask1 = bwareafilt(mask, mask_thd2);
    dilation_factor=str2num(get(handles.thd3,'string'));
   mask = bwareaopen(grayImage,mask_thd1);
    se = strel('square',dilation_factor);
  scar_dilated = imdilate(mask1,se);
 
   end
 
      
   scar_pix=nnz(scar_dilated);
    pixel_size=str2num(get(handles.pix_size,'String'));
    scarArea=pixel_size*pixel_size*scar_pix;

   set(handles.scar_pixel,'String',num2str(scar_pix));
   set(handles.scar_area_value,'String',num2str(scarArea));

 axes(handles.axes4_dilated_scar)
imshow(scar_dilated);
title("Masked Scar Region")
axis off

if get(handles.whole_pix_value,'String')==' '
    set(handles.scar_percent,'String', ' ');
else 
    scar_percent=scar_pix*100/str2num(get(handles.whole_pix_value,'String'));
    scar_percent=round(scar_percent,2);
    set(handles.scar_percent,'String',num2str(scar_percent));
end

handles.scar_dilated=scar_dilated;
handles.scar_pix=scar_pix;

handles.scarArea=scarArea;

guidata(hObject, handles);
function thd1_Callback(hObject, eventdata, handles)
% hObject    handle to thd1 (see GCBO)
% eventdata  reserved - to be defined in a future version of MATLAB
% handles    structure with handles and user data (see GUIDATA)

% Hints: get(hObject,'String') returns contents of thd1 as text
%        str2double(get(hObject,'String')) returns contents of thd1 as a double


% --- Executes during object creation, after setting all properties.
function thd1_CreateFcn(hObject, eventdata, handles)
% hObject    handle to thd1 (see GCBO)
% eventdata  reserved - to be defined in a future version of MATLAB
% handles    empty - handles not created until after all CreateFcns called

% Hint: edit controls usually have a white background on Windows.
%       See ISPC and COMPUTER.
if ispc && isequal(get(hObject,'BackgroundColor'), get(0,'defaultUicontrolBackgroundColor'))
    set(hObject,'BackgroundColor','white');
end


% --- Executes on button press in save_highlighted.
function save_highlighted_Callback(hObject, eventdata, handles)
% hObject    handle to save_highlighted (see GCBO)
% eventdata  reserved - to be defined in a future version of MATLAB
% handles    structure with handles and user data (see GUIDATA)
% imwrite(mask, [scar_folder, '\', filename], 'tiff', 'Compression', 'none');
flg_scar_folder=handles.flg_scar_folder;
bg_indicator=handles.bg_indicator;
if flg_scar_folder==0
    msgbox("A folder for saving images is needed. Please open/create scar folder");
end
if flg_scar_folder==1
scar_folder=handles.scar_folder;
fname_ori=handles.fname_ori;

    if rem(bg_indicator,2)==0
    img_highlighted_scar=handles.img_highlighted;
    elseif rem(bg_indicator,2)==1
    img_highlighted_scar=handles.img_highlighted_white;
    end
scar_imageRGB=strcat(fname_ori(1:end-4),'_highlighted.tif');
imwrite(img_highlighted_scar, [scar_folder, '\', scar_imageRGB], 'tif', 'Compression', 'none');
set(handles.save_highlighted,'ForegroundColor','b');
end
% --- Executes on button press in save_scar.
function save_scar_Callback(hObject, eventdata, handles)
% hObject    handle to save_scar (see GCBO)
% eventdata  reserved - to be defined in a future version of MATLAB
% handles    structure with handles and user data (see GUIDATA)
flg_scar_folder=handles.flg_scar_folder;
if flg_scar_folder==0
    msgbox("A folder for saving images is needed. Please open/create scar folder");
end
if flg_scar_folder==1
scar_folder=handles.scar_folder;
fname_ori=handles.fname_ori;
scar_cropped=handles.scar_cropped;
scar_imageRGB=strcat(fname_ori(1:end-4),'_cropped.tif');
imwrite(scar_cropped, [scar_folder, '\', scar_imageRGB], 'tif', 'Compression', 'none');   
set(handles.save_scar,'ForegroundColor','b');

end
% --- Executes on button press in save_dilated_scar.
function save_dilated_scar_Callback(hObject, eventdata, handles)
% hObject    handle to save_dilated_scar (see GCBO)
% eventdata  reserved - to be defined in a future version of MATLAB
% handles    structure with handles and user data (see GUIDATA)
flg_scar_folder=handles.flg_scar_folder;
if flg_scar_folder==0
    msgbox("A folder for saving images is needed. Please open/create scar folder");
end
if flg_scar_folder==1
scar_dilated=handles.scar_dilated;
scar_folder=handles.scar_folder;
fname_ori=handles.fname_ori;

scar_imageRGB=strcat(fname_ori(1:end-4),'_dilated.tif');
imwrite(scar_dilated, [scar_folder, '\', scar_imageRGB], 'tif', 'Compression', 'none');   
set(handles.save_dilated_scar,'ForegroundColor','b');

end
% --- Executes on button press in save_scar_info.
function save_scar_info_Callback(hObject, eventdata, handles)
% hObject    handle to save_scar_info (see GCBO)
% eventdata  reserved - to be defined in a future version of MATLAB
% handles    structure with handles and user data (see GUIDATA)
flg_data_folder=handles.flg_data_folder;
if flg_data_folder==0
    msgbox("A folder for saving data is needed. Please select a folder for results");
end
if flg_data_folder==1
results_folder=handles.results_folder;
filname=handles.fname_ori;
% scar_pix=handles.scar_pix;
% scarArea=handles.scarArea;
% whole_pix=handles.whole_pix;
% tissueArea=handles.tissueArea;

csvoutput = [results_folder, '\scar_area_results.csv'];
if exist(fullfile(results_folder, '\scar_area_results.csv')) == 0
    title_str=cat(2,'File Name',',','Pixel Size (mm)',',','Scar Pixels',',','Tissue Pixels',',','Scar percentage',',','Scar Area (mm^2)',',','Tissue Area (mm^2)');
    fid=fopen(csvoutput,'at+');
    fprintf(fid,title_str);
    fprintf(fid,'\n');
    fclose(fid);

%     dlmwrite(csvoutput, title_str, 'delimiter', ' ', '-append');
end
out=cat(2,filname,',', get(handles.pix_size,'String'),',',get(handles.scar_pixel,'String'),',',get(handles.whole_pix_value,'String'),',',get(handles.scar_percent,'String'),',',get(handles.scar_area_value,'String'),',',get(handles.tissue_area_value,'String'),',');
% dlmwrite(csvoutput, out, 'delimiter', ' ', '-append');
fid=fopen(csvoutput,'at+');
fprintf(fid,out);
fprintf(fid,'\n');
fclose(fid);
set(handles.save_scar_info,'ForegroundColor','b');
end

% --- Executes on button press in dilated_whole_tissue.
function dilated_whole_tissue_Callback(hObject, eventdata, handles)
% hObject    handle to dilated_whole_tissue (see GCBO)
% eventdata  reserved - to be defined in a future version of MATLAB
% handles    structure with handles and user data (see GUIDATA)
img_ori=handles.img_ori;
img_double_whole=im2double(img_ori);
% grayImage=imbinarize(img_double_whole);

[row,col,clr]=size(img_double_whole);
img_thd=zeros(row,col,clr);
bg_thd=str2num(get(handles.bg_thd_whole,'String'));


 if get(handles.he_staining,'Value')==1 
     %----------change background to black
    for b=1:1:row
        for c=1:1:col

        if img_double_whole(b,c,1)>bg_thd&&img_double_whole(b,c,2)>bg_thd&&img_double_whole(b,c,3)>bg_thd
               img_thd(b,c,1)=0;
               img_thd(b,c,2)=0;
               img_thd(b,c,3)=0;
        elseif img_double_whole(b,c,1)<0.2&&img_double_whole(b,c,2)<0.2&&img_double_whole(b,c,3)<0.2

                img_thd(b,c,1)=0;
                img_thd(b,c,2)=0;
                img_thd(b,c,3)=0;

        else
            img_thd(b,c,1)=img_double_whole(b,c,1);
            img_thd(b,c,2)=img_double_whole(b,c,2);
            img_thd(b,c,3)=img_double_whole(b,c,3);

        end  
        end
    end


 img_double_whole=img_thd;
 grayImage = 255*rgb2gray(img_double_whole);

mask_thd1=str2num(get(handles.size_whole_thd1,'string'));
  mask = grayImage>mask_thd1;

mask_thd2=str2num(get(handles.size_whole_thd2,'string'));


dilation_factor=str2num(get(handles.dilation_value_whole,'string'));
   se = strel('square',dilation_factor);
  img_dilation = imdilate(mask,se);

   whole_dilated = bwareafilt(img_dilation, mask_thd2);

 end

 if get(handles.psr_staining,'Value')==1 

 %----------change background to black
    for b=1:1:row
        for c=1:1:col

        if img_double_whole(b,c,1)>bg_thd&&img_double_whole(b,c,2)>bg_thd&&img_double_whole(b,c,3)>bg_thd
               img_thd(b,c,1)=0;
               img_thd(b,c,2)=0;
               img_thd(b,c,3)=0;
        elseif img_double_whole(b,c,1)<0.2&&img_double_whole(b,c,2)<0.2&&img_double_whole(b,c,3)<0.2

                img_thd(b,c,1)=0;
                img_thd(b,c,2)=0;
                img_thd(b,c,3)=0;

        else
            img_thd(b,c,1)=img_double_whole(b,c,1);
            img_thd(b,c,2)=img_double_whole(b,c,2);
            img_thd(b,c,3)=img_double_whole(b,c,3);

        end  
        end
    end


 img_double_whole=img_thd;
 grayImage = 255*rgb2gray(img_double_whole);

mask_thd1=str2num(get(handles.size_whole_thd1,'string'));
  mask = grayImage>mask_thd1;

mask_thd2=str2num(get(handles.size_whole_thd2,'string'));


dilation_factor=str2num(get(handles.dilation_value_whole,'string'));
   se = strel('square',dilation_factor);
  img_dilation = imdilate(mask,se);

   whole_dilated = bwareafilt(img_dilation, mask_thd2);

 end

if get(handles.mts_staining,'Value')==1 
 %----------change background to black
    for b=1:1:row
        for c=1:1:col

        if img_double_whole(b,c,1)>bg_thd&&img_double_whole(b,c,2)>bg_thd&&img_double_whole(b,c,3)>bg_thd
               img_thd(b,c,1)=0;
               img_thd(b,c,2)=0;
               img_thd(b,c,3)=0;
        elseif img_double_whole(b,c,1)<0.2&&img_double_whole(b,c,2)<0.2&&img_double_whole(b,c,3)<0.2

                img_thd(b,c,1)=0;
                img_thd(b,c,2)=0;
                img_thd(b,c,3)=0;

        else
            img_thd(b,c,1)=img_double_whole(b,c,1);
            img_thd(b,c,2)=img_double_whole(b,c,2);
            img_thd(b,c,3)=img_double_whole(b,c,3);

        end  
        end
    end


 img_double_whole=img_thd;
 grayImage = 255*rgb2gray(img_double_whole);
    
mask_thd1=str2num(get(handles.size_whole_thd1,'string'));
  mask = grayImage>mask_thd1;

mask_thd2=str2num(get(handles.size_whole_thd2,'string'));


dilation_factor=str2num(get(handles.dilation_value_whole,'string'));
   se = strel('square',dilation_factor);
  img_dilation = imdilate(mask,se);

   whole_dilated = bwareafilt(img_dilation, mask_thd2);

end

 if get(handles.ttc_staining,'Value')==1 

    for b=1:1:row
        for c=1:1:col

        if img_double_whole(b,c,1)>bg_thd||img_double_whole(b,c,2)>bg_thd||img_double_whole(b,c,3)>bg_thd
               img_thd(b,c,1)=0;
               img_thd(b,c,2)=0;
               img_thd(b,c,3)=0;
        else
            img_thd(b,c,1)=img_double_whole(b,c,1);
            img_thd(b,c,2)=img_double_whole(b,c,2);    
            img_thd(b,c,3)=img_double_whole(b,c,3);
        end
        end  
    end

%----remove weak signals
bg_thd_ttc=str2num(get(handles.bg_thd_ttc,'string'));

 for b=1:1:row
        for c=1:1:col

        if max(img_thd(b,c,:))<bg_thd_ttc
               img_thd(b,c,1)=0;
               img_thd(b,c,2)=0;
               img_thd(b,c,3)=0;
        
        end
        end  
 end

 for b=1:1:row
        for c=1:1:col

        if img_thd(b,c,3)>img_thd(b,c,1)+0.1
               img_thd(b,c,1)=0;
               img_thd(b,c,2)=0;
               img_thd(b,c,3)=0;
        
        end
        end  
 end

     
  img_double_whole=img_thd;
 grayImage = 255*rgb2gray(img_double_whole);

mask_thd1=str2num(get(handles.size_whole_thd1,'string'));
  mask = grayImage>mask_thd1;

mask_thd2=str2num(get(handles.size_whole_thd2,'string'));


dilation_factor=str2num(get(handles.dilation_value_whole,'string'));
   se = strel('square',dilation_factor);
  img_dilation = imdilate(mask,se);

   whole_dilated = bwareafilt(img_dilation, mask_thd2);

 end


   
     whole_pix=nnz(whole_dilated);
     pixel_size=str2num(get(handles.pix_size,'String')); 
     tissueArea=pixel_size*pixel_size*whole_pix;

axes(handles.axes6)
imshow(whole_dilated)
title('Masked whole tissue')
axis off
 set(handles.whole_pix_value,'String',num2str(whole_pix));
 set(handles.tissue_area_value,'String',num2str(tissueArea));

if get(handles.scar_pixel,'String')==' '
    set(handles.scar_percent,'String', ' ');
else 
    scar_percent=str2num(get(handles.scar_pixel,'String'))*100/whole_pix;
    scar_percent=round(scar_percent,2);
    set(handles.scar_percent,'String',num2str(scar_percent));
end

 handles.whole_pix=whole_pix;
 handles.tissueArea=tissueArea;
 handles.whole_dilated=whole_dilated;
 guidata(hObject, handles);

% --- Executes on button press in save_dilated_pixels.
function save_dilated_pixels_Callback(hObject, eventdata, handles)
% hObject    handle to save_dilated_pixels (see GCBO)
% eventdata  reserved - to be defined in a future version of MATLAB
% handles    structure with handles and user data (see GUIDATA)


% --- Executes on button press in scar_dir.
function scar_dir_Callback(hObject, eventdata, handles)
% hObject    handle to scar_dir (see GCBO)
% eventdata  reserved - to be defined in a future version of MATLAB
% handles    structure with handles and user data (see GUIDATA)
flg_scar_folder=handles.flg_scar_folder;
scar_folder=uigetdir;
flg_scar_folder=1;
index_scar_save=1;
handles.scar_folder=scar_folder;
handles.flg_scar_folder=flg_scar_folder;
guidata(hObject, handles);

% --- Executes on button press in data_dir.
function data_dir_Callback(hObject, eventdata, handles)
% hObject    handle to data_dir (see GCBO)
% eventdata  reserved - to be defined in a future version of MATLAB
% handles    structure with handles and user data (see GUIDATA)
flg_data_folder=handles.flg_data_folder;
results_folder=uigetdir;

flg_data_folder=1;
handles.results_folder=results_folder;
handles.flg_data_folder=flg_data_folder;
guidata(hObject, handles);

% --- Executes on button press in open_image.
function open_image_Callback(hObject, eventdata, handles)
% hObject    handle to open_image (see GCBO)
% eventdata  reserved - to be defined in a future version of MATLAB
% handles    structure with handles and user data (see GUIDATA)
[filname,pathname] = uigetfile({'*.tif';'*.png';'*.jpg';'*.gif'}, 'Select File to work with');
img_ori = imread([pathname, '\', filname]);

set(handles.file_name_display,'String',filname);
if get(handles.he_staining,'Value')==1
set(handles.color_thd,'String','0.7');
set(handles.thd1, 'String','50');
set(handles.thd2, 'String','50');
set(handles.thd3, 'String','20');
set(handles.bg_thd_whole,'String', '0.7');
set(handles.size_whole_thd1, 'String','10');
set(handles.size_whole_thd2, 'String','2');
set(handles.dilation_value_whole, 'String','20');

end
if get(handles.mts_staining,'Value')==1
set(handles.color_thd,'String','1.2');
set(handles.thd1, 'String','15');
set(handles.thd2, 'String','20');
set(handles.thd3, 'String','20');
end
if get(handles.ttc_staining,'Value')==1
set(handles.color_thd,'String','0.15');
set(handles.thd1, 'String','500');
set(handles.thd2, 'String','3');
set(handles.thd3, 'String','4');
end
if get(handles.psr_staining,'Value')==1
set(handles.color_thd,'String','0.15');
set(handles.thd1, 'String','5');
set(handles.thd2, 'String','20');
set(handles.thd3, 'String','3');
end


set(handles.scar_pixel,'String',' ');
set(handles.scar_area_value,'String',' ');
set(handles.whole_pix_value,'String',' ');
set(handles.tissue_area_value,'String',' ');
set(handles.scar_percent,'String',' ');
set(handles.save_highlighted,'ForegroundColor','black');
set(handles.save_extracted,'ForegroundColor','black');
set(handles.save_scar,'ForegroundColor','black');
set(handles.save_dilated_scar,'ForegroundColor','black');
set(handles.save_scar_info,'ForegroundColor','black');


cla(handles.axes2_highlighted);
cla(handles.axes2_highlighted2);
cla(handles.axes3_scar);
cla(handles.axes4_dilated_scar);
cla(handles.axes6);

reset(handles.axes2_highlighted);
reset(handles.axes2_highlighted2);
reset(handles.axes3_scar);
reset(handles.axes4_dilated_scar);
reset(handles.axes6);

%-------display the original image

axes(handles.axes1_original)

imshow(img_ori)
title('Original Image');
axis off

handles.img_ori=img_ori;
handles.fname_ori=filname;
handles.pname_ori=pathname;

guidata(hObject, handles);


% --- Executes on button press in scar_highlighted2.
function scar_highlighted2_Callback(hObject, eventdata, handles)
% hObject    handle to scar_highlighted2 (see GCBO)
% eventdata  reserved - to be defined in a future version of MATLAB
% handles    structure with handles and user data (see GUIDATA)
img_highlighted2=handles.img_highlighted;

row=handles.row_img;
col=handles.col_img;


if get(handles.he_staining,'Value')==1

    for b=1:1:row
      for c=1:1:col

%          end
            if img_highlighted2(b,c,1)>=img_highlighted2(b,c,2)||img_highlighted2(b,c,3)>=img_highlighted2(b,c,2)
                img_highlighted2(b,c,1)=0;
                img_highlighted2(b,c,2)=0;
                img_highlighted2(b,c,3)=0;

            end

      end
    end

h=fspecial('disk',5);
img_highlighted2=imfilter(img_highlighted2,h);

end

if get(handles.mts_staining,'Value')==1

    for b=1:1:row
      for c=1:1:col
%          if img_filtered_repeat(b,c,1)>clr_r_thd
%             img_filtered_repeat(b,c,1)=0;
%          end
%          if img_filtered_repeat(b,c,2)>clr_g_thd
%              img_filtered_repeat(b,c,2)=0;
%          end
%          if img_filtered_repeat(b,c,3)>clr_b_thd
%              img_filtered_repeat(b,c,3)=0;
%          end
            if img_highlighted2(b,c,1)>=img_highlighted2(b,c,3)||img_highlighted2(b,c,2)>=img_highlighted2(b,c,3)
                img_highlighted2(b,c,1)=0;
                img_highlighted2(b,c,2)=0;
                img_highlighted2(b,c,3)=0;
% %                 img_filtered_repeat(b,c,1)=1;
%                 img_filtered_repeat(b,c,2)=1;
% %                 img_filtered_repeat(b,c,3)=1;
            end
%             if img_filtered_repeat(b,c,2)<0.5
%                 img_filtered_repeat(b,c,2)=0;
%             end

      end
    end

h=fspecial('disk',1);
img_highlighted2=imfilter(img_highlighted2,h);

%  nColors = 32;
%  img_gray = rgb2ind(img_filtered_repeat,32);
% % 
% % se = strel('line',10,100);
% % img_gray = imerode(img_gray,se);
% % 
%  se1 = strel('line',50,100);
% 
%   img_gray= imdilate(img_gray,se1);
%--------------filter method 1------------
end

 axes(handles.axes2_highlighted2)
 imshow(img_highlighted2)
 title('Separated Scar')
 axis off


handles.img_scar=img_highlighted2;
 guidata(hObject, handles);


function thd2_Callback(hObject, eventdata, handles)
% hObject    handle to thd2 (see GCBO)
% eventdata  reserved - to be defined in a future version of MATLAB
% handles    structure with handles and user data (see GUIDATA)

% Hints: get(hObject,'String') returns contents of thd2 as text
%        str2double(get(hObject,'String')) returns contents of thd2 as a double


% --- Executes during object creation, after setting all properties.
function thd2_CreateFcn(hObject, eventdata, handles)
% hObject    handle to thd2 (see GCBO)
% eventdata  reserved - to be defined in a future version of MATLAB
% handles    empty - handles not created until after all CreateFcns called

% Hint: edit controls usually have a white background on Windows.
%       See ISPC and COMPUTER.
if ispc && isequal(get(hObject,'BackgroundColor'), get(0,'defaultUicontrolBackgroundColor'))
    set(hObject,'BackgroundColor','white');
end


% --- Executes on button press in overlap_scar.
function overlap_scar_Callback(hObject, eventdata, handles)
% hObject    handle to overlap_scar (see GCBO)
% eventdata  reserved - to be defined in a future version of MATLAB
% handles    structure with handles and user data (see GUIDATA)
scar_dilated=handles.scar_dilated;
scar_cropped=handles.scar_cropped;
scar_overlapped=imfuse(scar_cropped,scar_dilated,'falsecolor');
figure;
imshow(scar_overlapped);


function scar_pixel_Callback(hObject, eventdata, handles)
% hObject    handle to scar_pixel (see GCBO)
% eventdata  reserved - to be defined in a future version of MATLAB
% handles    structure with handles and user data (see GUIDATA)

% Hints: get(hObject,'String') returns contents of scar_pixel as text
%        str2double(get(hObject,'String')) returns contents of scar_pixel as a double


% --- Executes during object creation, after setting all properties.
function scar_pixel_CreateFcn(hObject, eventdata, handles)
% hObject    handle to scar_pixel (see GCBO)
% eventdata  reserved - to be defined in a future version of MATLAB
% handles    empty - handles not created until after all CreateFcns called

% Hint: edit controls usually have a white background on Windows.
%       See ISPC and COMPUTER.
if ispc && isequal(get(hObject,'BackgroundColor'), get(0,'defaultUicontrolBackgroundColor'))
    set(hObject,'BackgroundColor','white');
end


function pix_size_Callback(hObject, eventdata, handles)
% hObject    handle to pix_size (see GCBO)
% eventdata  reserved - to be defined in a future version of MATLAB
% handles    structure with handles and user data (see GUIDATA)

% Hints: get(hObject,'String') returns contents of pix_size as text
%        str2double(get(hObject,'String')) returns contents of pix_size as a double


% --- Executes during object creation, after setting all properties.
function pix_size_CreateFcn(hObject, eventdata, handles)
% hObject    handle to pix_size (see GCBO)
% eventdata  reserved - to be defined in a future version of MATLAB
% handles    empty - handles not created until after all CreateFcns called

% Hint: edit controls usually have a white background on Windows.
%       See ISPC and COMPUTER.
if ispc && isequal(get(hObject,'BackgroundColor'), get(0,'defaultUicontrolBackgroundColor'))
    set(hObject,'BackgroundColor','white');
end


function scar_area_value_Callback(hObject, eventdata, handles)
% hObject    handle to scar_area_value (see GCBO)
% eventdata  reserved - to be defined in a future version of MATLAB
% handles    structure with handles and user data (see GUIDATA)

% Hints: get(hObject,'String') returns contents of scar_area_value as text
%        str2double(get(hObject,'String')) returns contents of scar_area_value as a double


% --- Executes during object creation, after setting all properties.
function scar_area_value_CreateFcn(hObject, eventdata, handles)
% hObject    handle to scar_area_value (see GCBO)
% eventdata  reserved - to be defined in a future version of MATLAB
% handles    empty - handles not created until after all CreateFcns called

% Hint: edit controls usually have a white background on Windows.
%       See ISPC and COMPUTER.
if ispc && isequal(get(hObject,'BackgroundColor'), get(0,'defaultUicontrolBackgroundColor'))
    set(hObject,'BackgroundColor','white');
end


function file_name_display_Callback(hObject, eventdata, handles)
% hObject    handle to file_name_display (see GCBO)
% eventdata  reserved - to be defined in a future version of MATLAB
% handles    structure with handles and user data (see GUIDATA)

% Hints: get(hObject,'String') returns contents of file_name_display as text
%        str2double(get(hObject,'String')) returns contents of file_name_display as a double


% --- Executes during object creation, after setting all properties.
function file_name_display_CreateFcn(hObject, eventdata, handles)
% hObject    handle to file_name_display (see GCBO)
% eventdata  reserved - to be defined in a future version of MATLAB
% handles    empty - handles not created until after all CreateFcns called

% Hint: edit controls usually have a white background on Windows.
%       See ISPC and COMPUTER.
if ispc && isequal(get(hObject,'BackgroundColor'), get(0,'defaultUicontrolBackgroundColor'))
    set(hObject,'BackgroundColor','white');
end


% --- Executes on button press in he_staining.
function he_staining_Callback(hObject, eventdata, handles)
% hObject    handle to he_staining (see GCBO)
% eventdata  reserved - to be defined in a future version of MATLAB
% handles    structure with handles and user data (see GUIDATA)

% Hint: get(hObject,'Value') returns toggle state of he_staining
if get(handles.he_staining,'Value')==1
    set(handles.mts_staining,'Value',0);
    set(handles.he_staining,'ForegroundColor','b');
    set(handles.mts_staining,'ForegroundColor','k');
    set(handles.color_thd,'String',num2str(0.7));
    set(handles.thd1,'String',num2str(50));
    set(handles.thd2,'String',num2str(50));
    set(handles.thd3,'String',num2str(20));
    set(handles.ttc_staining,'Value',0);
    set(handles.ttc_staining,'ForegroundColor','k');
    set(handles.psr_staining,'Value',0);
    set(handles.psr_staining,'ForegroundColor','k');
    set(handles.bg_thd_whole,'String', '0.7');
    set(handles.size_whole_thd1, 'String','10');
    set(handles.size_whole_thd2, 'String','2');
    set(handles.dilation_value_whole, 'String','20');
    set(handles.bg_thd_ttc,'enable','off')
end

% --- Executes on button press in mts_staining.
function mts_staining_Callback(hObject, eventdata, handles)
% hObject    handle to mts_staining (see GCBO)
% eventdata  reserved - to be defined in a future version of MATLAB
% handles    structure with handles and user data (see GUIDATA)

% Hint: get(hObject,'Value') returns toggle state of mts_staining
if get(handles.mts_staining,'Value')==1
    set(handles.he_staining,'Value',0);
    set(handles.mts_staining,'ForegroundColor','b');
    set(handles.he_staining,'ForegroundColor','k');
    set(handles.color_thd,'String',num2str(1.2));
    set(handles.thd1,'String',num2str(15));
    set(handles.thd2,'String',num2str(20));
    
    set(handles.ttc_staining,'Value',0);
    set(handles.ttc_staining,'ForegroundColor','k');
    set(handles.psr_staining,'Value',0);
    set(handles.psr_staining,'ForegroundColor','k');
    set(handles.bg_thd_ttc,'enable','off')
end


function whole_pix_value_Callback(hObject, eventdata, handles)
% hObject    handle to whole_pix_value (see GCBO)
% eventdata  reserved - to be defined in a future version of MATLAB
% handles    structure with handles and user data (see GUIDATA)

% Hints: get(hObject,'String') returns contents of whole_pix_value as text
%        str2double(get(hObject,'String')) returns contents of whole_pix_value as a double


% --- Executes during object creation, after setting all properties.
function whole_pix_value_CreateFcn(hObject, eventdata, handles)
% hObject    handle to whole_pix_value (see GCBO)
% eventdata  reserved - to be defined in a future version of MATLAB
% handles    empty - handles not created until after all CreateFcns called

% Hint: edit controls usually have a white background on Windows.
%       See ISPC and COMPUTER.
if ispc && isequal(get(hObject,'BackgroundColor'), get(0,'defaultUicontrolBackgroundColor'))
    set(hObject,'BackgroundColor','white');
end


function tissue_area_value_Callback(hObject, eventdata, handles)
% hObject    handle to tissue_area_value (see GCBO)
% eventdata  reserved - to be defined in a future version of MATLAB
% handles    structure with handles and user data (see GUIDATA)

% Hints: get(hObject,'String') returns contents of tissue_area_value as text
%        str2double(get(hObject,'String')) returns contents of tissue_area_value as a double


% --- Executes during object creation, after setting all properties.
function tissue_area_value_CreateFcn(hObject, eventdata, handles)
% hObject    handle to tissue_area_value (see GCBO)
% eventdata  reserved - to be defined in a future version of MATLAB
% handles    empty - handles not created until after all CreateFcns called

% Hint: edit controls usually have a white background on Windows.
%       See ISPC and COMPUTER.
if ispc && isequal(get(hObject,'BackgroundColor'), get(0,'defaultUicontrolBackgroundColor'))
    set(hObject,'BackgroundColor','white');
end


% --- Executes on button press in save_masked_image.
function save_masked_image_Callback(hObject, eventdata, handles)
% hObject    handle to save_masked_image (see GCBO)
% eventdata  reserved - to be defined in a future version of MATLAB
% handles    structure with handles and user data (see GUIDATA)
flg_scar_folder=handles.flg_scar_folder;
if flg_scar_folder==0
    msgbox("A folder for saving images is needed. Please open/create scar folder");
end
if flg_scar_folder==1

whole_dilated=handles.whole_dilated;
scar_folder=handles.scar_folder;
fname_ori=handles.fname_ori;
whole_image_name=strcat(fname_ori(1:end-4),'_dilated_image.tif');
imwrite(whole_dilated, [scar_folder, '\', whole_image_name], 'tif', 'Compression', 'none');
set(handles.save_masked_image,'ForegroundColor','b');
end


function dilation_value_whole_Callback(hObject, eventdata, handles)
% hObject    handle to dilation_value_whole (see GCBO)
% eventdata  reserved - to be defined in a future version of MATLAB
% handles    structure with handles and user data (see GUIDATA)

% Hints: get(hObject,'String') returns contents of dilation_value_whole as text
%        str2double(get(hObject,'String')) returns contents of dilation_value_whole as a double


% --- Executes during object creation, after setting all properties.
function dilation_value_whole_CreateFcn(hObject, eventdata, handles)
% hObject    handle to dilation_value_whole (see GCBO)
% eventdata  reserved - to be defined in a future version of MATLAB
% handles    empty - handles not created until after all CreateFcns called

% Hint: edit controls usually have a white background on Windows.
%       See ISPC and COMPUTER.
if ispc && isequal(get(hObject,'BackgroundColor'), get(0,'defaultUicontrolBackgroundColor'))
    set(hObject,'BackgroundColor','white');
end


% --- Executes on button press in save_extracted.
function save_extracted_Callback(hObject, eventdata, handles)
% hObject    handle to save_extracted (see GCBO)
% eventdata  reserved - to be defined in a future version of MATLAB
% handles    structure with handles and user data (see GUIDATA)
scar_extracted=handles.img_scar;

flg_scar_folder=handles.flg_scar_folder;
if flg_scar_folder==0
    msgbox("A folder for saving images is needed. Please open/create scar folder");
end
if flg_scar_folder==1
scar_folder=handles.scar_folder;
fname_ori=handles.fname_ori;

scar_extracted=handles.img_scar;
scar_imageRGB=strcat(fname_ori(1:end-4),'_extracted.tif');
imwrite(scar_extracted, [scar_folder, '\', scar_imageRGB], 'tif', 'Compression', 'none');
set(handles.save_extracted,'ForegroundColor','b');
end


% --- Executes on button press in ttc_staining.
function ttc_staining_Callback(hObject, eventdata, handles)
% hObject    handle to ttc_staining (see GCBO)
% eventdata  reserved - to be defined in a future version of MATLAB
% handles    structure with handles and user data (see GUIDATA)

% Hint: get(hObject,'Value') returns toggle state of ttc_staining
if get(handles.ttc_staining,'Value')==1
    set(handles.he_staining,'Value',0);
    set(handles.ttc_staining,'ForegroundColor','b');
    set(handles.he_staining,'ForegroundColor','k');
%     set(handles.color_thd,'String',num2str(1.2));
%     set(handles.thd1,'String',num2str(15));
%     set(handles.thd2,'String',num2str(20));
    
    set(handles.mts_staining,'Value',0);
    set(handles.mts_staining,'ForegroundColor','k');
    set(handles.psr_staining,'Value',0);
    set(handles.psr_staining,'ForegroundColor','k');
    set(handles.color_thd,'String',num2str(0.15));
    set(handles.thd1,'String',num2str(500));
    set(handles.thd2,'String',num2str(3));
    set(handles.thd3,'String',num2str(4));
    set(handles.bg_thd_ttc,'enable','on')
   
end

% --- Executes on button press in psr_staining.
function psr_staining_Callback(hObject, eventdata, handles)
% hObject    handle to psr_staining (see GCBO)
% eventdata  reserved - to be defined in a future version of MATLAB
% handles    structure with handles and user data (see GUIDATA)

% Hint: get(hObject,'Value') returns toggle state of psr_staining
if get(handles.psr_staining,'Value')==1
    set(handles.he_staining,'Value',0);
    set(handles.psr_staining,'ForegroundColor','b');
    set(handles.he_staining,'ForegroundColor','k');
%     set(handles.color_thd,'String',num2str(1.2));
%     set(handles.thd1,'String',num2str(15));
%     set(handles.thd2,'String',num2str(20));
    
    set(handles.mts_staining,'Value',0);
    set(handles.mts_staining,'ForegroundColor','k');
    set(handles.ttc_staining,'Value',0);
    set(handles.ttc_staining,'ForegroundColor','k');
    set(handles.color_thd,'String',num2str(0.15));
    set(handles.thd1,'String',num2str(5));
    set(handles.thd2,'String',num2str(20));
    set(handles.thd3,'String',num2str(3));
    set(handles.bg_thd_whole,'String', '0.7');
    set(handles.size_whole_thd1, 'String','30');
    set(handles.size_whole_thd2, 'String','1');
    set(handles.dilation_value_whole, 'String','5');
    set(handles.bg_thd_ttc,'enable','off')
end


function thd3_Callback(hObject, eventdata, handles)
% hObject    handle to thd3 (see GCBO)
% eventdata  reserved - to be defined in a future version of MATLAB
% handles    structure with handles and user data (see GUIDATA)

% Hints: get(hObject,'String') returns contents of thd3 as text
%        str2double(get(hObject,'String')) returns contents of thd3 as a double


% --- Executes during object creation, after setting all properties.
function thd3_CreateFcn(hObject, eventdata, handles)
% hObject    handle to thd3 (see GCBO)
% eventdata  reserved - to be defined in a future version of MATLAB
% handles    empty - handles not created until after all CreateFcns called

% Hint: edit controls usually have a white background on Windows.
%       See ISPC and COMPUTER.
if ispc && isequal(get(hObject,'BackgroundColor'), get(0,'defaultUicontrolBackgroundColor'))
    set(hObject,'BackgroundColor','white');
end


function bg_thd_whole_Callback(hObject, eventdata, handles)
% hObject    handle to bg_thd_whole (see GCBO)
% eventdata  reserved - to be defined in a future version of MATLAB
% handles    structure with handles and user data (see GUIDATA)

% Hints: get(hObject,'String') returns contents of bg_thd_whole as text
%        str2double(get(hObject,'String')) returns contents of bg_thd_whole as a double


% --- Executes during object creation, after setting all properties.
function bg_thd_whole_CreateFcn(hObject, eventdata, handles)
% hObject    handle to bg_thd_whole (see GCBO)
% eventdata  reserved - to be defined in a future version of MATLAB
% handles    empty - handles not created until after all CreateFcns called

% Hint: edit controls usually have a white background on Windows.
%       See ISPC and COMPUTER.
if ispc && isequal(get(hObject,'BackgroundColor'), get(0,'defaultUicontrolBackgroundColor'))
    set(hObject,'BackgroundColor','white');
end


function size_whole_thd1_Callback(hObject, eventdata, handles)
% hObject    handle to size_whole_thd1 (see GCBO)
% eventdata  reserved - to be defined in a future version of MATLAB
% handles    structure with handles and user data (see GUIDATA)

% Hints: get(hObject,'String') returns contents of size_whole_thd1 as text
%        str2double(get(hObject,'String')) returns contents of size_whole_thd1 as a double


% --- Executes during object creation, after setting all properties.
function size_whole_thd1_CreateFcn(hObject, eventdata, handles)
% hObject    handle to size_whole_thd1 (see GCBO)
% eventdata  reserved - to be defined in a future version of MATLAB
% handles    empty - handles not created until after all CreateFcns called

% Hint: edit controls usually have a white background on Windows.
%       See ISPC and COMPUTER.
if ispc && isequal(get(hObject,'BackgroundColor'), get(0,'defaultUicontrolBackgroundColor'))
    set(hObject,'BackgroundColor','white');
end


function size_whole_thd2_Callback(hObject, eventdata, handles)
% hObject    handle to size_whole_thd2 (see GCBO)
% eventdata  reserved - to be defined in a future version of MATLAB
% handles    structure with handles and user data (see GUIDATA)

% Hints: get(hObject,'String') returns contents of size_whole_thd2 as text
%        str2double(get(hObject,'String')) returns contents of size_whole_thd2 as a double


% --- Executes during object creation, after setting all properties.
function size_whole_thd2_CreateFcn(hObject, eventdata, handles)
% hObject    handle to size_whole_thd2 (see GCBO)
% eventdata  reserved - to be defined in a future version of MATLAB
% handles    empty - handles not created until after all CreateFcns called

% Hint: edit controls usually have a white background on Windows.
%       See ISPC and COMPUTER.
if ispc && isequal(get(hObject,'BackgroundColor'), get(0,'defaultUicontrolBackgroundColor'))
    set(hObject,'BackgroundColor','white');
end


% --- Executes on button press in enlarge_dilated_whole.
function enlarge_dilated_whole_Callback(hObject, eventdata, handles)
% hObject    handle to enlarge_dilated_whole (see GCBO)
% eventdata  reserved - to be defined in a future version of MATLAB
% handles    structure with handles and user data (see GUIDATA)

img_dilated=handles.whole_dilated;
figure;
imshow(img_dilated)


function bg_thd_ttc_Callback(hObject, eventdata, handles)
% hObject    handle to bg_thd_ttc (see GCBO)
% eventdata  reserved - to be defined in a future version of MATLAB
% handles    structure with handles and user data (see GUIDATA)

% Hints: get(hObject,'String') returns contents of bg_thd_ttc as text
%        str2double(get(hObject,'String')) returns contents of bg_thd_ttc as a double


% --- Executes during object creation, after setting all properties.
function bg_thd_ttc_CreateFcn(hObject, eventdata, handles)
% hObject    handle to bg_thd_ttc (see GCBO)
% eventdata  reserved - to be defined in a future version of MATLAB
% handles    empty - handles not created until after all CreateFcns called

% Hint: edit controls usually have a white background on Windows.
%       See ISPC and COMPUTER.
if ispc && isequal(get(hObject,'BackgroundColor'), get(0,'defaultUicontrolBackgroundColor'))
    set(hObject,'BackgroundColor','white');
end


% --- Executes on button press in background_switch.
function background_switch_Callback(hObject, eventdata, handles)
% hObject    handle to background_switch (see GCBO)
% eventdata  reserved - to be defined in a future version of MATLAB
% handles    structure with handles and user data (see GUIDATA)

img_highlighted_white=handles.img_highlighted;
img_highlighted=handles.img_highlighted;
bg_indicator=handles.bg_indicator;
[row,col,clr]=size(img_highlighted_white);

if rem(bg_indicator,2)==0
    for b=1:1:row
      for c=1:1:col
         if img_highlighted_white(b,c,1)==0&&img_highlighted_white(b,c,2)==0&&img_highlighted_white(b,c,3)==0
             img_highlighted_white(b,c,1)=1;
             img_highlighted_white(b,c,2)=1;
             img_highlighted_white(b,c,3)=1;

         elseif img_highlighted_white(b,c,1)==0&&img_highlighted_white(b,c,2)==0&&img_highlighted_white(b,c,3)<0.2     
             img_highlighted_white(b,c,1)=1;
             img_highlighted_white(b,c,2)=1;
             img_highlighted_white(b,c,3)=1;
         end
      end
    end
    bg_indicator=bg_indicator+1;
   
   axes(handles.axes2_highlighted)
imshow(img_highlighted_white)
title('Image With High Contrast')
axis off 
handles.img_highlighted_white=img_highlighted_white;
elseif rem(bg_indicator,2)==1
    axes(handles.axes2_highlighted)
imshow(img_highlighted)
title('Image With High Contrast')
axis off 
bg_indicator=bg_indicator+1;
end

handles.bg_indicator=bg_indicator;
 guidata(hObject, handles);


function scar_percent_Callback(hObject, eventdata, handles)
% hObject    handle to scar_percent (see GCBO)
% eventdata  reserved - to be defined in a future version of MATLAB
% handles    structure with handles and user data (see GUIDATA)

% Hints: get(hObject,'String') returns contents of scar_percent as text
%        str2double(get(hObject,'String')) returns contents of scar_percent as a double


% --- Executes during object creation, after setting all properties.
function scar_percent_CreateFcn(hObject, eventdata, handles)
% hObject    handle to scar_percent (see GCBO)
% eventdata  reserved - to be defined in a future version of MATLAB
% handles    empty - handles not created until after all CreateFcns called

% Hint: edit controls usually have a white background on Windows.
%       See ISPC and COMPUTER.
if ispc && isequal(get(hObject,'BackgroundColor'), get(0,'defaultUicontrolBackgroundColor'))
    set(hObject,'BackgroundColor','white');
end


function pixels_highlight_Callback(hObject, eventdata, handles)
% hObject    handle to pixels_highlight (see GCBO)
% eventdata  reserved - to be defined in a future version of MATLAB
% handles    structure with handles and user data (see GUIDATA)

% Hints: get(hObject,'String') returns contents of pixels_highlight as text
%        str2double(get(hObject,'String')) returns contents of pixels_highlight as a double


% --- Executes during object creation, after setting all properties.
function pixels_highlight_CreateFcn(hObject, eventdata, handles)
% hObject    handle to pixels_highlight (see GCBO)
% eventdata  reserved - to be defined in a future version of MATLAB
% handles    empty - handles not created until after all CreateFcns called

% Hint: edit controls usually have a white background on Windows.
%       See ISPC and COMPUTER.
if ispc && isequal(get(hObject,'BackgroundColor'), get(0,'defaultUicontrolBackgroundColor'))
    set(hObject,'BackgroundColor','white');
end
```
