## Supplemental codes for "Detection, Isolation and Quantification of Myocardial Infarct with Four Different Histological Staining Techniques": split_image.m

```
function varargout = split_image(varargin)
% SPLIT_IMAGE MATLAB code for split_image.fig
%      SPLIT_IMAGE_MANUAL, by itself, creates a new SPLIT_IMAGE_MANUAL or raises the existing
%      singleton*.
%
%      H = SPLIT_IMAGE_MANUAL returns the handle to a new SPLIT_IMAGE_MANUAL or the handle to
%      the existing singleton*.
%
%      SPLIT_IMAGE_MANUAL('CALLBACK',hObject,eventData,handles,...) calls the local
%      function named CALLBACK in SPLIT_IMAGE_MANUAL.M with the given input arguments.
%
%      SPLIT_IMAGE_MANUAL('Property','Value',...) creates a new SPLIT_IMAGE_MANUAL or raises the
%      existing singleton*.  Starting from the left, property value pairs are
%      applied to the GUI before split_image_manual_OpeningFcn gets called.  An
%      unrecognized property name or invalid value makes property application
%      stop.  All inputs are passed to split_image_manual_OpeningFcn via varargin.
%
%      *See GUI Options on GUIDE's Tools menu.  Choose "GUI allows only one
%      instance to run (singleton)".
%
% See also: GUIDE, GUIDATA, GUIHANDLES

% Edit the above text to modify the response to help split_image_manual

% Last Modified by GUIDE v2.5 15-Jul-2024 12:21:01

% Begin initialization code - DO NOT EDIT
gui_Singleton = 1;
gui_State = struct('gui_Name',       mfilename, ...
                   'gui_Singleton',  gui_Singleton, ...
                   'gui_OpeningFcn', @split_image_OpeningFcn, ...
                   'gui_OutputFcn',  @split_image_OutputFcn, ...
                   'gui_LayoutFcn',  [] , ...
                   'gui_Callback',   []);
if nargin && ischar(varargin{1})
    gui_State.gui_Callback = str2func(varargin{1});
end

if nargout
    [varargout{1:nargout}] = gui_mainfcn(gui_State, varargin{:});
else
    gui_mainfcn(gui_State, varargin{:});
end
% End initialization code - DO NOT EDIT


% --- Executes just before split_image_manual is made visible.
function split_image_OpeningFcn(hObject, eventdata, handles, varargin)
% This function has no output args, see OutputFcn.
% hObject    handle to figure
% eventdata  reserved - to be defined in a future version of MATLAB
% handles    structure with handles and user data (see GUIDATA)
% varargin   command line arguments to split_image_manual (see VARARGIN)

% Choose default command line output for split_image_manual
handles.output = hObject;
set(handles.nImage_split,'string','3')
set(handles.top_pix_image_4,'enable','off')
set(handles.bottom_pix_image_4,'enable','off')
set(handles.top_pix_image_5,'enable','off')
set(handles.bottom_pix_image_5,'enable','off')

% Update handles structure
guidata(hObject, handles);

% UIWAIT makes split_image_manual wait for user response (see UIRESUME)
% uiwait(handles.figure1);


% --- Outputs from this function are returned to the command line.
function varargout = split_image_OutputFcn(hObject, eventdata, handles) 
% varargout  cell array for returning output args (see VARARGOUT);
% hObject    handle to figure
% eventdata  reserved - to be defined in a future version of MATLAB
% handles    structure with handles and user data (see GUIDATA)

% Get default command line output from handles structure
varargout{1} = handles.output;


% --- Executes on button press in image_1.
function image_1_Callback(hObject, eventdata, handles)
% hObject    handle to image_1 (see GCBO)
% eventdata  reserved - to be defined in a future version of MATLAB
% handles    structure with handles and user data (see GUIDATA)

% Hint: get(hObject,'Value') returns toggle state of image_1
if get(handles.image_1,'value')==1
    set(handles.image_2,'value',0);
    set(handles.image_3,'value',0);
    set(handles.image_4,'value',0);
    set(handles.image_5,'value',0);
    set(handles.image_6,'value',0);
    set(handles.top_pix_image_1,'string',' ');
    set(handles.bottom_pix_image_1,'string',' ');
end

% --- Executes on button press in image_2.
function image_2_Callback(hObject, eventdata, handles)
% hObject    handle to image_2 (see GCBO)
% eventdata  reserved - to be defined in a future version of MATLAB
% handles    structure with handles and user data (see GUIDATA)

% Hint: get(hObject,'Value') returns toggle state of image_2
if get(handles.image_2,'value')==1
    set(handles.image_1,'value',0);
    set(handles.image_3,'value',0);
    set(handles.image_4,'value',0);
    set(handles.image_5,'value',0);
    set(handles.image_6,'value',0);
    set(handles.top_pix_image_1,'string',' ');
    set(handles.bottom_pix_image_1,'string',' ');
end

% --- Executes on button press in image_3.
function image_3_Callback(hObject, eventdata, handles)
% hObject    handle to image_3 (see GCBO)
% eventdata  reserved - to be defined in a future version of MATLAB
% handles    structure with handles and user data (see GUIDATA)

% Hint: get(hObject,'Value') returns toggle state of image_3
if get(handles.image_3,'value')==1
    set(handles.image_2,'value',0);
    set(handles.image_1,'value',0);
    set(handles.image_4,'value',0);
    set(handles.image_5,'value',0);
    set(handles.image_6,'value',0);
    set(handles.top_pix_image_1,'string',' ');
    set(handles.bottom_pix_image_1,'string',' ');
end

% --- Executes on button press in image_4.
function image_4_Callback(hObject, eventdata, handles)
% hObject    handle to image_4 (see GCBO)
% eventdata  reserved - to be defined in a future version of MATLAB
% handles    structure with handles and user data (see GUIDATA)

% Hint: get(hObject,'Value') returns toggle state of image_4
if get(handles.image_4,'value')==1
    set(handles.image_2,'value',0);
    set(handles.image_3,'value',0);
    set(handles.image_1,'value',0);
    set(handles.image_5,'value',0);
    set(handles.image_6,'value',0);
    set(handles.top_pix_image_1,'string',' ');
    set(handles.bottom_pix_image_1,'string',' ');
end

% --- Executes on button press in image_5.
function image_5_Callback(hObject, eventdata, handles)
% hObject    handle to image_5 (see GCBO)
% eventdata  reserved - to be defined in a future version of MATLAB
% handles    structure with handles and user data (see GUIDATA)

% Hint: get(hObject,'Value') returns toggle state of image_5
if get(handles.image_5,'value')==1
    set(handles.image_2,'value',0);
    set(handles.image_3,'value',0);
    set(handles.image_4,'value',0);
    set(handles.image_1,'value',0);
    set(handles.image_6,'value',0);
    set(handles.top_pix_image_1,'string',' ');
    set(handles.bottom_pix_image_1,'string',' ');
end

% --- Executes on button press in image_6.
function image_6_Callback(hObject, eventdata, handles)
% hObject    handle to image_6 (see GCBO)
% eventdata  reserved - to be defined in a future version of MATLAB
% handles    structure with handles and user data (see GUIDATA)

% Hint: get(hObject,'Value') returns toggle state of image_6
if get(handles.image_6,'value')==1
    set(handles.image_2,'value',0);
    set(handles.image_3,'value',0);
    set(handles.image_4,'value',0);
    set(handles.image_5,'value',0);
    set(handles.image_1,'value',0);
    set(handles.top_pix_image_1,'string',' ');
    set(handles.bottom_pix_image_1,'string',' ');
end

% --- Executes on button press in split_image.
function split_image_Callback(hObject, eventdata, handles)
% hObject    handle to split_image (see GCBO)
% eventdata  reserved - to be defined in a future version of MATLAB
% handles    structure with handles and user data (see GUIDATA)
img_ori=handles.img_ori;
col=handles.col_img;
clr=handles.clr_img;
row=handles.row_img;
nimage=str2num(get(handles.nImage_split,'string'));

cla(handles.axes2);
cla(handles.axes4);
cla(handles.axes5);
cla(handles.axes6);
cla(handles.axes7);

if nimage==1

image_1=img_ori(str2num(get(handles.top_pix_image_1,'string')):str2num(get(handles.bottom_pix_image_1,'string')),:,:);

axes(handles.axes2)
imshow(image_1)
title('image 1')

handles.image_1=image_1;

end

if nimage==2

image_1=img_ori(str2num(get(handles.top_pix_image_1,'string')):str2num(get(handles.bottom_pix_image_1,'string')),:,:);
image_2=img_ori(str2num(get(handles.top_pix_image_2,'string'))+1:str2num(get(handles.bottom_pix_image_2,'string')),:,:);

axes(handles.axes2)
imshow(image_1)
title('image 1')
axes(handles.axes4)
imshow(image_2)
title('image 2')

handles.image_1=image_1;
handles.image_2=image_2;
end

if nimage==3

image_1=img_ori(str2num(get(handles.top_pix_image_1,'string')):str2num(get(handles.bottom_pix_image_1,'string')),:,:);
image_2=img_ori(str2num(get(handles.top_pix_image_2,'string'))+1:str2num(get(handles.bottom_pix_image_2,'string')),:,:);
image_3=img_ori(str2num(get(handles.top_pix_image_3,'string'))+1:str2num(get(handles.bottom_pix_image_3,'string')),:,:);

axes(handles.axes2)
imshow(image_1)
title('image 1')
axes(handles.axes4)
imshow(image_2)
title('image 2')

axes(handles.axes5)
imshow(image_3)
title('image 3')

handles.image_1=image_1;
handles.image_2=image_2;
handles.image_3=image_3;
end

if nimage==4

image_1=img_ori(str2num(get(handles.top_pix_image_1,'string')):str2num(get(handles.bottom_pix_image_1,'string')),:,:);
image_2=img_ori(str2num(get(handles.top_pix_image_2,'string')):str2num(get(handles.bottom_pix_image_2,'string')),:,:);
image_3=img_ori(str2num(get(handles.top_pix_image_3,'string')):str2num(get(handles.bottom_pix_image_3,'string')),:,:);
image_4=img_ori(str2num(get(handles.top_pix_image_4,'string')):str2num(get(handles.bottom_pix_image_4,'string')),:,:);

axes(handles.axes2)
imshow(image_1)
title('image 1')
axes(handles.axes4)
imshow(image_2)
title('image 2')
axes(handles.axes5)
imshow(image_3)
title('image 3')
axes(handles.axes6)
imshow(image_4)
title('image 4')

handles.image_1=image_1;
handles.image_2=image_2;
handles.image_3=image_3;
handles.image_4=image_4;
end

if nimage==5

image_1=img_ori(str2num(get(handles.top_pix_image_1,'string')):str2num(get(handles.bottom_pix_image_1,'string')),:,:);
image_2=img_ori(str2num(get(handles.top_pix_image_2,'string')):str2num(get(handles.bottom_pix_image_2,'string')),:,:);
image_3=img_ori(str2num(get(handles.top_pix_image_3,'string')):str2num(get(handles.bottom_pix_image_3,'string')),:,:);
image_4=img_ori(str2num(get(handles.top_pix_image_4,'string')):str2num(get(handles.bottom_pix_image_4,'string')),:,:);
image_5=img_ori(str2num(get(handles.top_pix_image_5,'string')):str2num(get(handles.bottom_pix_image_5,'string')),:,:);
axes(handles.axes2)
imshow(image_1)
title('image 1')
axes(handles.axes4)
imshow(image_2)
title('image 2')
axes(handles.axes5)
imshow(image_3)
title('image 3')
axes(handles.axes6)
imshow(image_4)
title('image 4')
axes(handles.axes7)
imshow(image_5)
title('image 5')
handles.image_1=image_1;
handles.image_2=image_2;
handles.image_3=image_3;
handles.image_4=image_4;
handles.image_5=image_5;

end


guidata(hObject, handles);
function thd_split_Callback(hObject, eventdata, handles)
% hObject    handle to thd_split (see GCBO)
% eventdata  reserved - to be defined in a future version of MATLAB
% handles    structure with handles and user data (see GUIDATA)

% Hints: get(hObject,'String') returns contents of thd_split as text
%        str2double(get(hObject,'String')) returns contents of thd_split as a double


% --- Executes during object creation, after setting all properties.
function thd_split_CreateFcn(hObject, eventdata, handles)
% hObject    handle to thd_split (see GCBO)
% eventdata  reserved - to be defined in a future version of MATLAB
% handles    empty - handles not created until after all CreateFcns called

% Hint: edit controls usually have a white background on Windows.
%       See ISPC and COMPUTER.
if ispc && isequal(get(hObject,'BackgroundColor'), get(0,'defaultUicontrolBackgroundColor'))
    set(hObject,'BackgroundColor','white');
end


function filname_Callback(hObject, eventdata, handles)
% hObject    handle to filname (see GCBO)
% eventdata  reserved - to be defined in a future version of MATLAB
% handles    structure with handles and user data (see GUIDATA)

% Hints: get(hObject,'String') returns contents of filname as text
%        str2double(get(hObject,'String')) returns contents of filname as a double


% --- Executes during object creation, after setting all properties.
function filname_CreateFcn(hObject, eventdata, handles)
% hObject    handle to filname (see GCBO)
% eventdata  reserved - to be defined in a future version of MATLAB
% handles    empty - handles not created until after all CreateFcns called

% Hint: edit controls usually have a white background on Windows.
%       See ISPC and COMPUTER.
if ispc && isequal(get(hObject,'BackgroundColor'), get(0,'defaultUicontrolBackgroundColor'))
    set(hObject,'BackgroundColor','white');
end


function top_pix_image_1_Callback(hObject, eventdata, handles)
% hObject    handle to top_pix_image_1 (see GCBO)
% eventdata  reserved - to be defined in a future version of MATLAB
% handles    structure with handles and user data (see GUIDATA)

% Hints: get(hObject,'String') returns contents of top_pix_image_1 as text
%        str2double(get(hObject,'String')) returns contents of top_pix_image_1 as a double


% --- Executes during object creation, after setting all properties.
function top_pix_image_1_CreateFcn(hObject, eventdata, handles)
% hObject    handle to top_pix_image_1 (see GCBO)
% eventdata  reserved - to be defined in a future version of MATLAB
% handles    empty - handles not created until after all CreateFcns called

% Hint: edit controls usually have a white background on Windows.
%       See ISPC and COMPUTER.
if ispc && isequal(get(hObject,'BackgroundColor'), get(0,'defaultUicontrolBackgroundColor'))
    set(hObject,'BackgroundColor','white');
end


function bottom_pix_image_1_Callback(hObject, eventdata, handles)
% hObject    handle to bottom_pix_image_1 (see GCBO)
% eventdata  reserved - to be defined in a future version of MATLAB
% handles    structure with handles and user data (see GUIDATA)

% Hints: get(hObject,'String') returns contents of bottom_pix_image_1 as text
%        str2double(get(hObject,'String')) returns contents of bottom_pix_image_1 as a double


% --- Executes during object creation, after setting all properties.
function bottom_pix_image_1_CreateFcn(hObject, eventdata, handles)
% hObject    handle to bottom_pix_image_1 (see GCBO)
% eventdata  reserved - to be defined in a future version of MATLAB
% handles    empty - handles not created until after all CreateFcns called

% Hint: edit controls usually have a white background on Windows.
%       See ISPC and COMPUTER.
if ispc && isequal(get(hObject,'BackgroundColor'), get(0,'defaultUicontrolBackgroundColor'))
    set(hObject,'BackgroundColor','white');
end


% --- Executes on button press in save_image.
function save_image_Callback(hObject, eventdata, handles)
% hObject    handle to save_image (see GCBO)
% eventdata  reserved - to be defined in a future version of MATLAB
% handles    structure with handles and user data (see GUIDATA)

filname=handles.filename;
nimage=str2num(get(handles.nImage_split,'string'));
if nimage==1
image_1=handles.image_1;

image_1_name=strcat(filname(1:end-4),'_img1.tif');

pathname_save=uigetdir(matlabroot,'MATLAB Root Folder');
  
imwrite(image_1, [pathname_save, '\', image_1_name], 'tif', 'Compression', 'none');
msgbox('Image is saved');

end
if nimage==2
image_1=handles.image_1;
image_2=handles.image_2;

image_1_name=strcat(filname(1:end-4),'_img1.tif');
image_2_name=strcat(filname(1:end-4),'_img2.tif');

pathname_save=uigetdir(matlabroot,'MATLAB Root Folder');
  
imwrite(image_1, [pathname_save, '\', image_1_name], 'tif', 'Compression', 'none');
imwrite(image_2, [pathname_save, '\', image_2_name], 'tif', 'Compression', 'none');
msgbox('Image is saved');

end

if nimage==3
image_1=handles.image_1;
image_2=handles.image_2;
image_3=handles.image_3;

image_1_name=strcat(filname(1:end-4),'_img1.tif');
image_2_name=strcat(filname(1:end-4),'_img2.tif');
image_3_name=strcat(filname(1:end-4),'_img3.tif');

pathname_save=uigetdir(matlabroot,'MATLAB Root Folder');
  
imwrite(image_1, [pathname_save, '\', image_1_name], 'tif', 'Compression', 'none');
imwrite(image_2, [pathname_save, '\', image_2_name], 'tif', 'Compression', 'none');
imwrite(image_3, [pathname_save, '\', image_3_name], 'tif', 'Compression', 'none');
msgbox('Image is saved');

end

if nimage==4
image_1=handles.image_1;
image_2=handles.image_2;
image_3=handles.image_3;
image_4=handles.image_4;

image_1_name=strcat(filname(1:end-4),'_img1.tif');
image_2_name=strcat(filname(1:end-4),'_img2.tif');
image_3_name=strcat(filname(1:end-4),'_img3.tif');
image_4_name=strcat(filname(1:end-4),'_img4.tif');

pathname_save=uigetdir(matlabroot,'MATLAB Root Folder');
  
imwrite(image_1, [pathname_save, '\', image_1_name], 'tif', 'Compression', 'none');
imwrite(image_2, [pathname_save, '\', image_2_name], 'tif', 'Compression', 'none');
imwrite(image_3, [pathname_save, '\', image_3_name], 'tif', 'Compression', 'none');
imwrite(image_4, [pathname_save, '\', image_4_name], 'tif', 'Compression', 'none');
msgbox('Image is saved');

end

if nimage==5
image_1=handles.image_1;
image_2=handles.image_2;
image_3=handles.image_3;
image_4=handles.image_4;
image_5=handles.image_5;

image_1_name=strcat(filname(1:end-4),'_img1.tif');
image_2_name=strcat(filname(1:end-4),'_img2.tif');
image_3_name=strcat(filname(1:end-4),'_img3.tif');
image_4_name=strcat(filname(1:end-4),'_img4.tif');
image_5_name=strcat(filname(1:end-4),'_img5.tif');

pathname_save=uigetdir(matlabroot,'MATLAB Root Folder');
  
imwrite(image_1, [pathname_save, '\', image_1_name], 'tif', 'Compression', 'none');
imwrite(image_2, [pathname_save, '\', image_2_name], 'tif', 'Compression', 'none');
imwrite(image_3, [pathname_save, '\', image_3_name], 'tif', 'Compression', 'none');
imwrite(image_4, [pathname_save, '\', image_4_name], 'tif', 'Compression', 'none');
imwrite(image_5, [pathname_save, '\', image_5_name], 'tif', 'Compression', 'none');
msgbox('Image is saved');

end
% image_folder=handles.pathname;
% 
% if get(handles.image_1,'value')==1
% fn_image=strcat(filname(1:end-4),'_img1.tif');
% end
% if get(handles.image_2,'value')==1
% fn_image=strcat(filname(1:end-4),'_img2.tif');
% end
% if get(handles.image_3,'value')==1
% fn_image=strcat(filname(1:end-4),'_img3.tif');
% end
% if get(handles.image_4,'value')==1
% fn_image=strcat(filname(1:end-4),'_img4.tif');
% end
% if get(handles.image_5,'value')==1
% fn_image=strcat(filname(1:end-4),'_img5.tif');
% end
% if get(handles.image_6,'value')==1
% fn_image=strcat(filname(1:end-4),'_img6.tif');
% end
% 
% imwrite(image_selc, [image_folder, '\', fn_image], 'tif', 'Compression', 'none');
% msgbox('Image is saved');

% --------------------------------------------------------------------
function file_menu_Callback(hObject, eventdata, handles)
% hObject    handle to file_menu (see GCBO)
% eventdata  reserved - to be defined in a future version of MATLAB
% handles    structure with handles and user data (see GUIDATA)


% --------------------------------------------------------------------
function load_image_Callback(hObject, eventdata, handles)
% hObject    handle to load_image (see GCBO)
% eventdata  reserved - to be defined in a future version of MATLAB
% handles    structure with handles and user data (see GUIDATA)

[filname,pathname] = uigetfile({'*.tif';'*.jpg';'*.png'}, 'Select File to work with');
set(handles.filname,'string',filname);
img_ori = imread([pathname, '\', filname]);

% img_double=im2double(img_ori);
% imshow(img_ori)

[row,col,clr]=size(img_ori);
% for a_color=0:0.05:1

axes(handles.axes1)
imshow(img_ori)
axis on

set(handles.nImage_split,'string','3');
set(handles.top_pix_image_1,'enable','on');
set(handles.bottom_pix_image_1,'enable','on'); 
set(handles.top_pix_image_2,'enable','on');
set(handles.bottom_pix_image_2,'enable','on');
set(handles.top_pix_image_3,'enable','on');
set(handles.bottom_pix_image_3,'enable','on');
set(handles.top_pix_image_4,'enable','off','string', ' ');
set(handles.bottom_pix_image_4,'enable','off','string',' ');
set(handles.top_pix_image_5,'enable','off','string',' ');
set(handles.bottom_pix_image_5,'enable','off','string',' ');


length_section=round(row/3);
set(handles.top_pix_image_1,'string',num2str(1));
set(handles.bottom_pix_image_1,'string',num2str(length_section));
set(handles.top_pix_image_2,'string',num2str(length_section+1));
set(handles.bottom_pix_image_2,'string',num2str(length_section*2));
set(handles.top_pix_image_3,'string',num2str(length_section*2+1));
set(handles.bottom_pix_image_3,'string',row);

axes(handles.axes1)
hold on
h1=plot([1,col],[1,1],'Color','r','LineWidth',2);
h2=plot([1,col],[length_section,length_section],'Color','r','LineWidth',2);
h3=plot([1,col],[length_section+10,length_section+10],'Color','g','LineWidth',2);
h4=plot([1,col],[length_section*2,length_section*2],'Color','g','LineWidth',2);
h5=plot([1,col],[length_section*2+10,length_section*2+10],'Color','b','LineWidth',2);
h6=plot([1,col],[row,row],'Color','b','LineWidth',2);

hold off


% handles.img_double=img_double;
handles.img_ori=img_ori;
handles.filename=filname;
handles.pathname=pathname;
handles.row_img=row;
handles.col_img=col;
handles.clr_img=clr;
guidata(hObject, handles);


function row_pix_Callback(hObject, eventdata, handles)
% hObject    handle to row_pix (see GCBO)
% eventdata  reserved - to be defined in a future version of MATLAB
% handles    structure with handles and user data (see GUIDATA)

% Hints: get(hObject,'String') returns contents of row_pix as text
%        str2double(get(hObject,'String')) returns contents of row_pix as a double


% --- Executes during object creation, after setting all properties.
function row_pix_CreateFcn(hObject, eventdata, handles)
% hObject    handle to row_pix (see GCBO)
% eventdata  reserved - to be defined in a future version of MATLAB
% handles    empty - handles not created until after all CreateFcns called

% Hint: edit controls usually have a white background on Windows.
%       See ISPC and COMPUTER.
if ispc && isequal(get(hObject,'BackgroundColor'), get(0,'defaultUicontrolBackgroundColor'))
    set(hObject,'BackgroundColor','white');
end


function column_pix_Callback(hObject, eventdata, handles)
% hObject    handle to column_pix (see GCBO)
% eventdata  reserved - to be defined in a future version of MATLAB
% handles    structure with handles and user data (see GUIDATA)

% Hints: get(hObject,'String') returns contents of column_pix as text
%        str2double(get(hObject,'String')) returns contents of column_pix as a double


% --- Executes during object creation, after setting all properties.
function column_pix_CreateFcn(hObject, eventdata, handles)
% hObject    handle to column_pix (see GCBO)
% eventdata  reserved - to be defined in a future version of MATLAB
% handles    empty - handles not created until after all CreateFcns called

% Hint: edit controls usually have a white background on Windows.
%       See ISPC and COMPUTER.
if ispc && isequal(get(hObject,'BackgroundColor'), get(0,'defaultUicontrolBackgroundColor'))
    set(hObject,'BackgroundColor','white');
end


function nImage_split_Callback(hObject, eventdata, handles)
% hObject    handle to nImage_split (see GCBO)
% eventdata  reserved - to be defined in a future version of MATLAB
% handles    structure with handles and user data (see GUIDATA)

% Hints: get(hObject,'String') returns contents of nImage_split as text
%        str2double(get(hObject,'String')) returns contents of nImage_split as a double


% --- Executes during object creation, after setting all properties.
function nImage_split_CreateFcn(hObject, eventdata, handles)
% hObject    handle to nImage_split (see GCBO)
% eventdata  reserved - to be defined in a future version of MATLAB
% handles    empty - handles not created until after all CreateFcns called

% Hint: edit controls usually have a white background on Windows.
%       See ISPC and COMPUTER.
if ispc && isequal(get(hObject,'BackgroundColor'), get(0,'defaultUicontrolBackgroundColor'))
    set(hObject,'BackgroundColor','white');
end


function bottom_pix_image_2_Callback(hObject, eventdata, handles)
% hObject    handle to bottom_pix_image_2 (see GCBO)
% eventdata  reserved - to be defined in a future version of MATLAB
% handles    structure with handles and user data (see GUIDATA)

% Hints: get(hObject,'String') returns contents of bottom_pix_image_2 as text
%        str2double(get(hObject,'String')) returns contents of bottom_pix_image_2 as a double


% --- Executes during object creation, after setting all properties.
function bottom_pix_image_2_CreateFcn(hObject, eventdata, handles)
% hObject    handle to bottom_pix_image_2 (see GCBO)
% eventdata  reserved - to be defined in a future version of MATLAB
% handles    empty - handles not created until after all CreateFcns called

% Hint: edit controls usually have a white background on Windows.
%       See ISPC and COMPUTER.
if ispc && isequal(get(hObject,'BackgroundColor'), get(0,'defaultUicontrolBackgroundColor'))
    set(hObject,'BackgroundColor','white');
end


function top_pix_image_2_Callback(hObject, eventdata, handles)
% hObject    handle to top_pix_image_2 (see GCBO)
% eventdata  reserved - to be defined in a future version of MATLAB
% handles    structure with handles and user data (see GUIDATA)

% Hints: get(hObject,'String') returns contents of top_pix_image_2 as text
%        str2double(get(hObject,'String')) returns contents of top_pix_image_2 as a double


% --- Executes during object creation, after setting all properties.
function top_pix_image_2_CreateFcn(hObject, eventdata, handles)
% hObject    handle to top_pix_image_2 (see GCBO)
% eventdata  reserved - to be defined in a future version of MATLAB
% handles    empty - handles not created until after all CreateFcns called

% Hint: edit controls usually have a white background on Windows.
%       See ISPC and COMPUTER.
if ispc && isequal(get(hObject,'BackgroundColor'), get(0,'defaultUicontrolBackgroundColor'))
    set(hObject,'BackgroundColor','white');
end


function bottom_pix_image_3_Callback(hObject, eventdata, handles)
% hObject    handle to bottom_pix_image_3 (see GCBO)
% eventdata  reserved - to be defined in a future version of MATLAB
% handles    structure with handles and user data (see GUIDATA)

% Hints: get(hObject,'String') returns contents of bottom_pix_image_3 as text
%        str2double(get(hObject,'String')) returns contents of bottom_pix_image_3 as a double


% --- Executes during object creation, after setting all properties.
function bottom_pix_image_3_CreateFcn(hObject, eventdata, handles)
% hObject    handle to bottom_pix_image_3 (see GCBO)
% eventdata  reserved - to be defined in a future version of MATLAB
% handles    empty - handles not created until after all CreateFcns called

% Hint: edit controls usually have a white background on Windows.
%       See ISPC and COMPUTER.
if ispc && isequal(get(hObject,'BackgroundColor'), get(0,'defaultUicontrolBackgroundColor'))
    set(hObject,'BackgroundColor','white');
end


function top_pix_image_3_Callback(hObject, eventdata, handles)
% hObject    handle to top_pix_image_3 (see GCBO)
% eventdata  reserved - to be defined in a future version of MATLAB
% handles    structure with handles and user data (see GUIDATA)

% Hints: get(hObject,'String') returns contents of top_pix_image_3 as text
%        str2double(get(hObject,'String')) returns contents of top_pix_image_3 as a double


% --- Executes during object creation, after setting all properties.
function top_pix_image_3_CreateFcn(hObject, eventdata, handles)
% hObject    handle to top_pix_image_3 (see GCBO)
% eventdata  reserved - to be defined in a future version of MATLAB
% handles    empty - handles not created until after all CreateFcns called

% Hint: edit controls usually have a white background on Windows.
%       See ISPC and COMPUTER.
if ispc && isequal(get(hObject,'BackgroundColor'), get(0,'defaultUicontrolBackgroundColor'))
    set(hObject,'BackgroundColor','white');
end


% --- Executes on button press in nImages_ok.
function nImages_ok_Callback(hObject, eventdata, handles)
% hObject    handle to nImages_ok (see GCBO)
% eventdata  reserved - to be defined in a future version of MATLAB
% handles    structure with handles and user data (see GUIDATA)
row=handles.row_img;
col=handles.col_img;
n_image=str2num(get(handles.nImage_split,'string'));
hplots=zeros(10,1);

if n_image==1

axesHandlesToChildObjects = findobj(handles.axes1, 'Type', 'line');
  if ~isempty(axesHandlesToChildObjects)
    delete(axesHandlesToChildObjects);
  end 
set(handles.top_pix_image_1,'enable','on');
set(handles.bottom_pix_image_1,'enable','on');   
set(handles.top_pix_image_2,'enable','off','string', ' ');
set(handles.bottom_pix_image_2,'enable','off','string', ' ');
set(handles.top_pix_image_3,'enable','off','string', ' ');
set(handles.bottom_pix_image_3,'enable','off','string', ' ');
set(handles.top_pix_image_4,'enable','off','string', ' ');
set(handles.bottom_pix_image_4,'enable','off','string', ' ');
set(handles.top_pix_image_5,'enable','off','string', ' ');
set(handles.bottom_pix_image_5,'enable','off','string', ' ');

length_section=round(row/n_image);
set(handles.top_pix_image_1,'string',num2str(1));
set(handles.bottom_pix_image_1,'string',num2str(length_section));

axes(handles.axes1)
hold on
h1=plot([1,col],[1,1],'Color','r','LineWidth',2);
h2=plot([1,col],[row,row],'Color','r','LineWidth',2);

hold off
hplots(1)=h1;
hplots(2)=h2;
end

if n_image==2

axesHandlesToChildObjects = findobj(handles.axes1, 'Type', 'line');
  if ~isempty(axesHandlesToChildObjects)
    delete(axesHandlesToChildObjects);
  end 

 set(handles.top_pix_image_1,'enable','on');
set(handles.bottom_pix_image_1,'enable','on'); 
    set(handles.top_pix_image_2,'enable','on');
set(handles.bottom_pix_image_2,'enable','on');
set(handles.top_pix_image_3,'enable','off','string', ' ');
set(handles.bottom_pix_image_3,'enable','off','string', ' ');
set(handles.top_pix_image_4,'enable','off','string', ' ');
set(handles.bottom_pix_image_4,'enable','off','string', ' ');
set(handles.top_pix_image_5,'enable','off','string', ' ');
set(handles.bottom_pix_image_5,'enable','off','string', ' ');

length_section=round(row/n_image);
set(handles.top_pix_image_1,'string',num2str(1));
set(handles.bottom_pix_image_1,'string',num2str(length_section));
set(handles.top_pix_image_2,'string',num2str(length_section+1));
set(handles.bottom_pix_image_2,'string',num2str(row));

axes(handles.axes1)
hold on
h1=plot([1,col],[1,1],'Color','r','LineWidth',2);
h2=plot([1,col],[length_section,length_section],'Color','r','LineWidth',2);
h3=plot([1,col],[length_section+10,length_section+10],'Color','g','LineWidth',2);
h4=plot([1,col],[row,row],'Color','g','LineWidth',2);

hold off
hplots(1)=h1;
hplots(2)=h2;
hplots(3)=h3;
hplots(4)=h4;
% h1_obj=gobjects(h1);
% handles.h1=h1_obj;
end
if n_image==3

    axesHandlesToChildObjects = findobj(handles.axes1, 'Type', 'line');
  if ~isempty(axesHandlesToChildObjects)
    delete(axesHandlesToChildObjects);
  end 

    set(handles.top_pix_image_1,'enable','on');
set(handles.bottom_pix_image_1,'enable','on'); 
 set(handles.top_pix_image_2,'enable','on');
set(handles.bottom_pix_image_2,'enable','on');
set(handles.top_pix_image_3,'enable','on');
set(handles.bottom_pix_image_3,'enable','on');
set(handles.top_pix_image_4,'enable','off','string', ' ');
set(handles.bottom_pix_image_4,'enable','off','string', ' ');
set(handles.top_pix_image_5,'enable','off','string', ' ');
set(handles.bottom_pix_image_5,'enable','off','string', ' ');

length_section=round(row/n_image);
set(handles.top_pix_image_1,'string',num2str(1));
set(handles.bottom_pix_image_1,'string',num2str(length_section));
set(handles.top_pix_image_2,'string',num2str(length_section+1));
set(handles.bottom_pix_image_2,'string',num2str(length_section*2));
set(handles.top_pix_image_3,'string',num2str(length_section*2+1));
set(handles.bottom_pix_image_3,'string',num2str(row));

axes(handles.axes1)
hold on
h1=plot([1,col],[1,1],'Color','r','LineWidth',2);
h2=plot([1,col],[length_section,length_section],'Color','r','LineWidth',2);
h3=plot([1,col],[length_section+10,length_section+10],'Color','g','LineWidth',2);
h4=plot([1,col],[length_section*2,length_section*2],'Color','g','LineWidth',2);
h5=plot([1,col],[length_section*2+10,length_section*2+10],'Color','b','LineWidth',2);
h6=plot([1,col],[row,row],'Color','b','LineWidth',2);

hold off
hplots(1)=h1;
hplots(2)=h2;
hplots(3)=h3;
hplots(4)=h4;
hplots(5)=h5;
hplots(6)=h6;
% h1_obj=gobjects(h1);
% h2_obj=gobjects(h2);
% handles.h1=h1_obj;
% handles.h2=h2_obj;

end
if n_image==4

axesHandlesToChildObjects = findobj(handles.axes1, 'Type', 'line');
  if ~isempty(axesHandlesToChildObjects)
    delete(axesHandlesToChildObjects);
  end 

    set(handles.top_pix_image_1,'enable','on');
set(handles.bottom_pix_image_1,'enable','on'); 
 set(handles.top_pix_image_2,'enable','on');
set(handles.bottom_pix_image_2,'enable','on');
set(handles.top_pix_image_3,'enable','on');
set(handles.bottom_pix_image_3,'enable','on');
set(handles.top_pix_image_4,'enable','on');
set(handles.bottom_pix_image_4,'enable','on');
set(handles.top_pix_image_5,'enable','off','string', ' ');
set(handles.bottom_pix_image_5,'enable','off','string', ' ');

length_section=round(row/n_image);
set(handles.top_pix_image_1,'string',num2str(1));
set(handles.bottom_pix_image_1,'string',num2str(length_section));
set(handles.top_pix_image_2,'string',num2str(length_section+1));
set(handles.bottom_pix_image_2,'string',num2str(length_section*2));
set(handles.top_pix_image_3,'string',num2str(length_section*2+1));
set(handles.bottom_pix_image_3,'string',num2str(length_section*3));
set(handles.top_pix_image_4,'string',num2str(length_section*3+1));
set(handles.bottom_pix_image_4,'string',num2str(row));

axes(handles.axes1)
hold on
h1=plot([1,col],[1,1],'Color','r','LineWidth',2);
h2=plot([1,col],[length_section,length_section],'Color','r','LineWidth',2);
h3=plot([1,col],[length_section+10,length_section+10],'Color','g','LineWidth',2);
h4=plot([1,col],[length_section*2,length_section*2],'Color','g','LineWidth',2);
h5=plot([1,col],[length_section*2+10,length_section*2+10],'Color','b','LineWidth',2);
h6=plot([1,col],[length_section*3,length_section*3],'Color','b','LineWidth',2);
h7=plot([1,col],[length_section*3+10,length_section*3+10],'Color','m','LineWidth',2);
h8=plot([1,col],[row,row],'Color','m','LineWidth',2);

hold off
hplots(1)=h1;
hplots(2)=h2;
hplots(3)=h3;
hplots(4)=h4;
hplots(5)=h5;
hplots(6)=h6;
hplots(7)=h7;
hplots(8)=h8;

% h1_obj=gobjects(h1);
% h2_obj=gobjects(h2);
% h3_obj=gobjects(h3);
% 
% handles.h1=h1_obj;
% handles.h2=h2_obj;
% handles.h3=h3_obj;

end
if n_image==5

axesHandlesToChildObjects = findobj(handles.axes1, 'Type', 'line');
  if ~isempty(axesHandlesToChildObjects)
    delete(axesHandlesToChildObjects);
  end 
    set(handles.top_pix_image_1,'enable','on');
set(handles.bottom_pix_image_1,'enable','on'); 
 set(handles.top_pix_image_2,'enable','on');
set(handles.bottom_pix_image_2,'enable','on');
set(handles.top_pix_image_3,'enable','on');
set(handles.bottom_pix_image_3,'enable','on');
set(handles.top_pix_image_4,'enable','on');
set(handles.bottom_pix_image_4,'enable','on');
set(handles.top_pix_image_5,'enable','on');
set(handles.bottom_pix_image_5,'enable','on');

length_section=round(row/n_image);
set(handles.top_pix_image_1,'string',num2str(1));
set(handles.bottom_pix_image_1,'string',num2str(length_section));
set(handles.top_pix_image_2,'string',num2str(length_section+1));
set(handles.bottom_pix_image_2,'string',num2str(length_section*2));
set(handles.top_pix_image_3,'string',num2str(length_section*2+1));
set(handles.bottom_pix_image_3,'string',num2str(length_section*3));
set(handles.top_pix_image_4,'string',num2str(length_section*3+1));
set(handles.bottom_pix_image_4,'string',num2str(length_section*4));
set(handles.top_pix_image_5,'string',num2str(length_section*4+1));
set(handles.bottom_pix_image_5,'string',num2str(row));

axes(handles.axes1)
hold on
h1=plot([1,col],[1,1],'Color','r','LineWidth',2);
h2=plot([1,col],[length_section,length_section],'Color','r','LineWidth',2);
h3=plot([1,col],[length_section+10,length_section+10],'Color','g','LineWidth',2);
h4=plot([1,col],[length_section*2,length_section*2],'Color','g','LineWidth',2);
h5=plot([1,col],[length_section*2+10,length_section*2+10],'Color','b','LineWidth',2);
h6=plot([1,col],[length_section*3,length_section*3],'Color','b','LineWidth',2);
h7=plot([1,col],[length_section*3+10,length_section*3+10],'Color','m','LineWidth',2);
h8=plot([1,col],[length_section*4,length_section*4],'Color','m','LineWidth',2);
h9=plot([1,col],[length_section*4+10,length_section*4+10],'Color','k','LineWidth',2);
h10=plot([1,col],[row,row],'Color','k','LineWidth',2);

hold off
hplots(1)=h1;
hplots(2)=h2;
hplots(3)=h3;
hplots(4)=h4;
hplots(5)=h5;
hplots(6)=h6;
hplots(7)=h7;
hplots(8)=h8;
hplots(9)=h9;
hplots(10)=h10;

% handles.h1=h1;
% handles.h2=h2;
% handles.h3=h3;
% handles.h4=h4;

end
handles.hplots=hplots;
guidata(hObject, handles);

function bottom_pix_image_5_Callback(hObject, eventdata, handles)
% hObject    handle to bottom_pix_image_5 (see GCBO)
% eventdata  reserved - to be defined in a future version of MATLAB
% handles    structure with handles and user data (see GUIDATA)

% Hints: get(hObject,'String') returns contents of bottom_pix_image_5 as text
%        str2double(get(hObject,'String')) returns contents of bottom_pix_image_5 as a double


% --- Executes during object creation, after setting all properties.
function bottom_pix_image_5_CreateFcn(hObject, eventdata, handles)
% hObject    handle to bottom_pix_image_5 (see GCBO)
% eventdata  reserved - to be defined in a future version of MATLAB
% handles    empty - handles not created until after all CreateFcns called

% Hint: edit controls usually have a white background on Windows.
%       See ISPC and COMPUTER.
if ispc && isequal(get(hObject,'BackgroundColor'), get(0,'defaultUicontrolBackgroundColor'))
    set(hObject,'BackgroundColor','white');
end


function top_pix_image_5_Callback(hObject, eventdata, handles)
% hObject    handle to top_pix_image_5 (see GCBO)
% eventdata  reserved - to be defined in a future version of MATLAB
% handles    structure with handles and user data (see GUIDATA)

% Hints: get(hObject,'String') returns contents of top_pix_image_5 as text
%        str2double(get(hObject,'String')) returns contents of top_pix_image_5 as a double


% --- Executes during object creation, after setting all properties.
function top_pix_image_5_CreateFcn(hObject, eventdata, handles)
% hObject    handle to top_pix_image_5 (see GCBO)
% eventdata  reserved - to be defined in a future version of MATLAB
% handles    empty - handles not created until after all CreateFcns called

% Hint: edit controls usually have a white background on Windows.
%       See ISPC and COMPUTER.
if ispc && isequal(get(hObject,'BackgroundColor'), get(0,'defaultUicontrolBackgroundColor'))
    set(hObject,'BackgroundColor','white');
end


function top_pix_image_4_Callback(hObject, eventdata, handles)
% hObject    handle to top_pix_image_4 (see GCBO)
% eventdata  reserved - to be defined in a future version of MATLAB
% handles    structure with handles and user data (see GUIDATA)

% Hints: get(hObject,'String') returns contents of top_pix_image_4 as text
%        str2double(get(hObject,'String')) returns contents of top_pix_image_4 as a double


% --- Executes during object creation, after setting all properties.
function top_pix_image_4_CreateFcn(hObject, eventdata, handles)
% hObject    handle to top_pix_image_4 (see GCBO)
% eventdata  reserved - to be defined in a future version of MATLAB
% handles    empty - handles not created until after all CreateFcns called

% Hint: edit controls usually have a white background on Windows.
%       See ISPC and COMPUTER.
if ispc && isequal(get(hObject,'BackgroundColor'), get(0,'defaultUicontrolBackgroundColor'))
    set(hObject,'BackgroundColor','white');
end


function bottom_pix_image_4_Callback(hObject, eventdata, handles)
% hObject    handle to bottom_pix_image_4 (see GCBO)
% eventdata  reserved - to be defined in a future version of MATLAB
% handles    structure with handles and user data (see GUIDATA)

% Hints: get(hObject,'String') returns contents of bottom_pix_image_4 as text
%        str2double(get(hObject,'String')) returns contents of bottom_pix_image_4 as a double


% --- Executes during object creation, after setting all properties.
function bottom_pix_image_4_CreateFcn(hObject, eventdata, handles)
% hObject    handle to bottom_pix_image_4 (see GCBO)
% eventdata  reserved - to be defined in a future version of MATLAB
% handles    empty - handles not created until after all CreateFcns called

% Hint: edit controls usually have a white background on Windows.
%       See ISPC and COMPUTER.
if ispc && isequal(get(hObject,'BackgroundColor'), get(0,'defaultUicontrolBackgroundColor'))
    set(hObject,'BackgroundColor','white');
end


% --- Executes on button press in cf_5.
function cf_5_Callback(hObject, eventdata, handles)
% hObject    handle to cf_5 (see GCBO)
% eventdata  reserved - to be defined in a future version of MATLAB
% handles    structure with handles and user data (see GUIDATA)
hplots=handles.hplots;
col=handles.col_img;
top_img5=str2num(get(handles.top_pix_image_5,'string'));
bottom_img5=str2num(get(handles.bottom_pix_image_5,'string'));
delete(hplots(9));
delete(hplots(10));
axes(handles.axes1)
hold on
h9=plot([1,col],[top_img5,top_img5],'Color','k','LineWidth',2);
h10=plot([1,col],[bottom_img5,bottom_img5],'Color','k','LineWidth',2);

hold off
hplots(9)=h9;
hplots(10)=h10;

handles.hplots=hplots;
guidata(hObject, handles);

% --- Executes on button press in cf_4.
function cf_4_Callback(hObject, eventdata, handles)
% hObject    handle to cf_4 (see GCBO)
% eventdata  reserved - to be defined in a future version of MATLAB
% handles    structure with handles and user data (see GUIDATA)
hplots=handles.hplots;
col=handles.col_img;
top_img4=str2num(get(handles.top_pix_image_4,'string'));
bottom_img4=str2num(get(handles.bottom_pix_image_4,'string'));
delete(hplots(7));
delete(hplots(8));
axes(handles.axes1)
hold on
h7=plot([1,col],[top_img4,top_img4],'Color','m','LineWidth',2);
h8=plot([1,col],[bottom_img4,bottom_img4],'Color','m','LineWidth',2);

hold off
hplots(7)=h7;
hplots(8)=h8;

handles.hplots=hplots;
guidata(hObject, handles);

% --- Executes on button press in cf_3.
function cf_3_Callback(hObject, eventdata, handles)
% hObject    handle to cf_3 (see GCBO)
% eventdata  reserved - to be defined in a future version of MATLAB
% handles    structure with handles and user data (see GUIDATA)
hplots=handles.hplots;
col=handles.col_img;
top_img3=str2num(get(handles.top_pix_image_3,'string'));
bottom_img3=str2num(get(handles.bottom_pix_image_3,'string'));
delete(hplots(5));
delete(hplots(6));
axes(handles.axes1)
hold on
h5=plot([1,col],[top_img3,top_img3],'Color','b','LineWidth',2);
h6=plot([1,col],[bottom_img3,bottom_img3],'Color','b','LineWidth',2);

hold off
hplots(5)=h5;
hplots(6)=h6;

handles.hplots=hplots;
guidata(hObject, handles);

% --- Executes on button press in cf_2.
function cf_2_Callback(hObject, eventdata, handles)
% hObject    handle to cf_2 (see GCBO)
% eventdata  reserved - to be defined in a future version of MATLAB
% handles    structure with handles and user data (see GUIDATA)
hplots=handles.hplots;
col=handles.col_img;
top_img2=str2num(get(handles.top_pix_image_2,'string'));
bottom_img2=str2num(get(handles.bottom_pix_image_2,'string'));
delete(hplots(3));
delete(hplots(4));
axes(handles.axes1)
hold on
h3=plot([1,col],[top_img2,top_img2],'Color','g','LineWidth',2);
h4=plot([1,col],[bottom_img2,bottom_img2],'Color','g','LineWidth',2);

hold off
hplots(3)=h3;
hplots(4)=h4;

handles.hplots=hplots;
guidata(hObject, handles);

% --- Executes on button press in cf_1.
function cf_1_Callback(hObject, eventdata, handles)
% hObject    handle to cf_1 (see GCBO)
% eventdata  reserved - to be defined in a future version of MATLAB
% handles    structure with handles and user data (see GUIDATA)
hplots=handles.hplots;
col=handles.col_img;
top_img1=str2num(get(handles.top_pix_image_1,'string'));
bottom_img1=str2num(get(handles.bottom_pix_image_1,'string'));
delete(hplots(1));
delete(hplots(2));
axes(handles.axes1)
hold on
h1=plot([1,col],[top_img1,top_img1],'Color','r','LineWidth',2);
h2=plot([1,col],[bottom_img1,bottom_img1],'Color','r','LineWidth',2);

hold off
hplots(1)=h1;
hplots(2)=h2;

handles.hplots=hplots;
guidata(hObject, handles);


% --- Executes on button press in rotate_image.
function rotate_image_Callback(hObject, eventdata, handles)
% hObject    handle to rotate_image (see GCBO)
% eventdata  reserved - to be defined in a future version of MATLAB
% handles    structure with handles and user data (see GUIDATA)
img_ori=handles.img_ori;

img_ori=imrotate(img_ori, 90);

axes(handles.axes1)
imshow(img_ori)
axis on

handles.img_ori=img_ori;
guidata(hObject, handles);
```
